# Evolutionarily conserved fMRI network dynamics in the mouse, macaque, and human brain

**DOI:** 10.1101/2023.07.19.549681

**Authors:** Daniel Gutierrez-Barragan, Julian S.B. Ramirez, Stefano Panzeri, Ting Xu, Alessandro Gozzi

**Affiliations:** Functional Neuroimaging Lab, Istituto Italiano di Tecnologia, Center for Neuroscience and Cognitive Systems, Rovereto, Italy; Center for the Developing Brain. Child Mind Institute, NY, USA; Institute for Neural Information Processing, Center for Molecular Neurobiology (ZMNH), University Medical Center Hamburg-Eppendorf (UKE), Hamburg, Germany

**Author notes:** Share senior authorship (Ting Xu, Alessandro Gozzi). Lead Author: Alessandro Gozzi PhD, Functional neuroimaging Laboratory, Istituto Italiano di Tecnologia, Center for Neuroscience and Cognitive Systems @ UNITN, Corso Bettini 31, 38068, Rovereto, Italy.

## Abstract

Evolutionarily relevant networks have been previously described in several mammalian species using time-averaged analyses of fMRI time-series. However, fMRI network activity is highly dynamic and continually evolves over timescales of seconds. Whether the dynamic organization of resting-state fMRI network activity is conserved across mammalian species remains unclear. Using frame-wise clustering of fMRI time-series, we find that intrinsic fMRI network dynamics in awake macaques and humans is characterized by recurrent transitions between a set of 4 dominant, neuroanatomically homologous fMRI coactivation modes (C-modes), three of which are also plausibly represented in the rodent brain. Importantly, in all species C-modes exhibit species-invariant dynamic features, including preferred occurrence at specific phases of fMRI global signal fluctuations, and a state transition structure compatible with infraslow coupled oscillator dynamics. Moreover, dominant C-mode occurrence reconstitutes the static organization of the fMRI connectome in all species, and is predictive of ranking of corresponding fMRI connectivity gradients. These results reveal a set of species-invariant principles underlying the dynamic organization of fMRI networks in mammalian species, and offer novel opportunities to relate fMRI network findings across the phylogenetic tree.

## INTRODUCTION

Spontaneous fluctuations in resting fMRI signals have been consistently shown to be temporally synchronized across multiple functional systems, delineating a set of reproducible topographies often referred to as Resting State Networks (RSNs)^1,2^. RSN mapping typically entails the computation of time-averaged statistical dependencies between fMRI time-series under the assumption that temporal structure of this activity is time-invariant over a time window of minutes^3^. Prompted by the need to complement human fMRI research with mechanistic investigations in physiologically accessible species^4,5^, multiple research group have begun to implement fMRI mapping in non-human primates and rodents^6,7^. These investigations have revealed interesting evolutionary correspondences in the organization of RSN across mammalian species. These encompass the presence of highly synchronous interhemispheric networks, including evolutionarily-relevant precursors of distributed integrative systems, such as the default mode (DMN) and salience networks^8–10^. However, spontaneous brain activity is highly dynamic and continuously evolves over the timescale of minutes^11,12^. Accordingly, a large body of experimental and theoretical work has shown that the correlation structure of RSNs varies across time^13,14^ and involves transient interactions between distinct functional systems that are continually revisited^15^. These observations suggest that mere time-invariant descriptions of spontaneous fMRI activity are not sufficient to comprehensively describe the functional architecture of the resting brain.

Although many approaches to study the dynamic organization of RSNs have been proposed^13,14^, frame-wise methods^16–19^ have recently gained traction as a flexible approach to investigate the dynamic organization of intrinsic fMRI activity. Compared to correlation-based approaches (e.g. sliding-window analyses), frame-wise approaches offer the possibility to (a) temporally localize the peaks and troughs of activity that underlie fMRI network dynamics and relate them to global fluctuations in brain activity; (b) describe the dynamic organization of fMRI using physiologically interpretable parameters (e.g. mean BOLD activity); and (c) identify the relevant dimensions of fMRI dynamics without the need to pre-impose regional parcellations. Using whole-brain framewise clustering of fMRI time-series to obtain coactivation patterns (CAPs), it has been recently shown that intrinsic fMRI activity is dominated by recurring, cyclic fluctuations between stereotypic functional topographies^16–19^. The simplicity of the CAP framework, its direct association with a directly quantifiable physiological property, as well as its high temporal and spatial resolution are perfectly suited to parsimoniously, yet comprehensively describe the dynamic organization of spontaneous network activity.

While previous studies have compared the static organization of fMRI networks across species^9,20^, attempts to directly relate the dynamic organization of intrinsic brain activity across the mammalian phylogenetic tree are lacking. Is fMRI network dynamics underpinned by a unifying set of species-invariant principles, or does this phenomenon instead reflect unique, species-specific attributes? And are fMRI dynamic states and their cyclic dynamics evolutionarily conserved, or do they encompass phylogenetically divergent motifs?

To address these questions, we leveraged fMRI datasets acquired in awake humans, macaques and mice to probe and compare the dynamic organization of fMRI in the mammalian brain. We find that fMRI network dynamics in all probed species is similarly characterized by cyclic transitions between a few dominant and neuroanatomically related fMRI “coactivation modes” (C-modes) which exhibit largely conserved topographies and dynamic features such as a quasi-periodic infraslow evolution and a structure of transitions between states compatible with a coupled-oscillators dynamics^21^. We further relate the occurrence of dynamic states represented by C-modes to the organization of the static connectome and fMRI connectivity gradients. These results suggest that resting fMRI activity in mammalian species is underpinned by evolutionarily conserved dynamic principles.

## RESULTS

### Dominant coactivation modes parsimoniously describe fMRI network dynamics in humans, macaques and mice

To compare the dynamic organization of fMRI network dynamics within an evolutionary perspective, we used a frame-wise approach based on the identification of coactivation patterns (CAPs)^17,22^. Data for this study consisted of awake rsfMRI datasets from two human cohorts (The Hangzhou Normal University -HNU: 30 subjects, 10 test-retest sessions^23^ and The Midnight Scan Club - MSC: 10 subjects, 5 test-retest sessions^24^); the Newcastle (NC) macaque cohort^25,26^ (8 animals with 2 test-retest sessions); and 44 mice under head-fixed conditions. Using k-means, we clustered fMRI frames in the concatenated timeseries given their spatial similarity. We averaged the frames in each cluster to produce group-level CAP maps^15–17^, then T-scored them at the voxel level. We then obtained single-subject CAP maps by averaging the corresponding clustered frames from each subject (**Figure S1**). To identify reproducible functional CAPs that are representative of the dynamic structure of fMRI networks in each species, we used a multi-criteria approach (see methods section) aimed at maximizing CAP reproducibility across individuals, sessions or datasets. This approach yielded k = 8 CAPs in human and macaques, and k = 6 in mice as optimal clustering solutions (see methods section and **Figures S2 and S3**). Previous work^16^ has shown that CAPs embody rich fMRI topographies that can be reliably matched into mirrored coactive and anti-coactive pairs characterized by opposite BOLD polarity (**Figure 1**). As we will demonstrate below, and similar to what has been observed in quasi-periodic patterns^27^,each matched “CAP and anti-CAP” pairs describe a cyclical fluctuation of a single fMRI state. As predicted, the employed procedure identified in all species mirrored CAP pairs characterized by opposite BOLD polarity (spatial correlation, r < -0.65, all species, all CAP pairs, **Figure 1A-B**). Importantly, the identified CAPs explained in all species a large proportion of variance in fMRI timeseries (R^2^ > 0.58 mice, >0.68 macaque, and >0.75 human). Moreover, they also guaranteed robust within and between dataset spatial and occurrence rate reproducibility (**Figures S2-S3**, permutation tests with random CAP identity shuffling). These results suggest that that a few “dominant” dynamic patterns can parsimoniously describe the dynamic organization of fMRI activity in multiple mammalian species.

**Figure 1.**
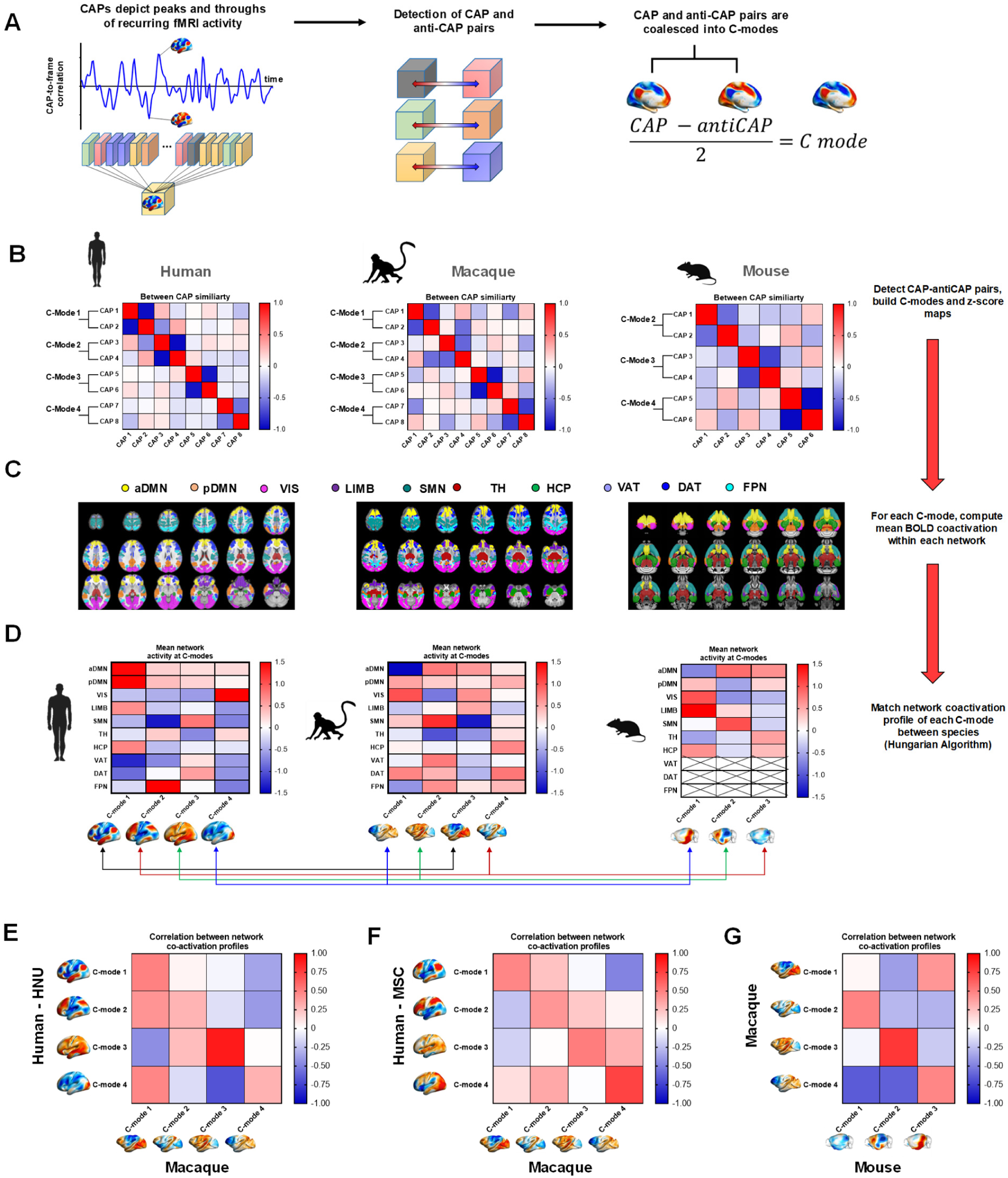
C-mode identification and matching across species. **(A)** CAPs represent transients of infraslow fMRI activity that can be matched in pairs exhibiting opposite polarity (CAPs, anti-CAPs, left). Detection of CAP anti-CAP pairs (middle) allows for the computation of fMRI Coactivation modes (right). **(B)** Detection of CAP-anti-CAP pairs given highest spatial anticorrelation in the between-CAP similarity matrix. C-modes are built by taking the highest occurring CAP from each pair, and spatially averaging it with its corresponding inverted anti-CAP. **(C)** Evolutionarily relevant fMRI networks used for matching. **(D)** Vectorized network coactivation profiles for each C-mode (spatially z-scored), extracted from the mean BOLD values of voxels within a network mask. Arrows denote matching of C-modes across species performed by the Hungarian Algorithm. **(E)** Correlation between matched C-modes from humans (HNU dataset) and macaques, **(F)** between humans (MSC dataset) and macaques; and **(G)** between macaques and mice. Abbreviations: aDMN - anterior Default Mode; pDMN - Posterior Default Mode; SMN – Somatomotor; VIS – Visual; DAT - Dorsal Attention; VAT-Ventral Attention; FPN – Frontoparietal; LIMB – Limbic; TH –Thalamus; HC – Hippocampus.

To further reduce dimensionality and facilitate cross-species comparisons, we coalesced highly anticorrelated CAP pairs into a single Coactivation Mode (C-mode). CAP and anti-CAP are thus the peak and trough of the same fluctuating C-mode. C-modes are thus computed by reversing the sign of the anti-CAP, and spatially averaging it with its paired CAP such to spatially depict the corresponding coactivation axis (**Figure 1A**). The topography of the individual CAPs pairs that constitute each C-mode is reported in **Figure S4**. To assess the potential confounding contribution of head-motion to C-mode topography mapping, we repeated the clustering procedure on time-series in which we did not scrub fMRI frames exhibiting high motion-related. We next compared the spatial topography of C-modes obtained with and without frame censoring. This comparison revealed that C-mode obtained using frame-censored timeseries were very similar to those we mapped using the entire timeseries (r > 0.95, all C-modes, all species, **Figure S5A**). Further corroborating a negligible contribution of head motion to our findings, we also found that no C-mode was preferentially enriched with high-motion frames in all species (one-way ANOVA, p > 0.3, F < 1.14 for all comparisons, **Figure S5B**).

### fMRI C-modes exhibit evolutionarily-conserved functional organization

Having identified dominant fMRI C-modes in all three species, we next asked whether their functional organization would show, on top of foreseeable species-specific features, recognizable evolutionarily conserved anatomic features. For this purpose, we matched C-mode topographies based on the similarity of mean BOLD coactivation profile across a set of evolutionarily conserved resting state networks (RSNs)^28–31^ (**Figure 1C**). The chosen networks include the Default-Mode (DMN), Visual (VIS), Somato-motor (SMN), Limbic (LIMB), Ventral (VAT) and Dorsal (DAT) Attention, Fronto-parietal (FPN), as well as key subcortical nuclei of the Thalamus (TH) and Hippocampus (HCP) and were selected based on the notion that, across these three species, they encompass partly-conserved neuroanatomical substrates^5,8^. A key exception of note is the lack of established phylogenetic precursors of the VAT, DAT, and FPN in the rodent brain^8^. For this reason, these networks were not included in the coactivation profile in mouse data. To allow spatial comparisons between species, C-mode maps were z-scored spatially and the mean of the normalized activity of voxels within each RSN mask was computed to build its corresponding profile vector (**Figure 1D**, C-modes numbered by decreasing occurrence rate). Vectors were first matched using the Hungarian Algorithm^32^ from human C-modes (organized in decreasing occurrence rate) to macaques, then from macaques to mice.

The corresponding results are depicted as spatial correlation between network coactivation profiles of human and macaque datasets (HNU and MSC, **Figure 1E and 1F**, respectively), as well as between macaques and mice (**Figure 1G**). Human to macaque matching gave consistent results for both HNU and MSC datasets, with the spatial of topography of human C-modes 1-4 being best aligned to corresponding macaque C-modes 1-4 (see also **Figure 2C**). Because the human HNU dataset included more subjects and was performed at daylight hours as animal scans, we describe our results hereafter for the HNU as main dataset and present a summary of the obtained MSC results as supplementary figure (**Figure S6**). Macaque to mouse matching linked macaque C-modes 2-4 to mouse C-mode 1-3, respectively (**Figure 1G**, **Figure 2C**). Whilst some anticorrelation was apparent in C-mode matching between single pairs of macaque and mouse C-modes, the chosen algorithmic solution ensured the overall best network matching across species (**Figure 2C**), as well as cross-species preservation of C-mode 3 and 4 phase coupling with fMRI global signal cycles (described below, cf. **Figure 4**). As human and macaque C-mode 1 was not preferentially matched to any mouse C-modes, we refer to mouse C-modes as 2, 3, and 4 for consistency with those mapped in higher species throughout the manuscript.

**Figure 2.**
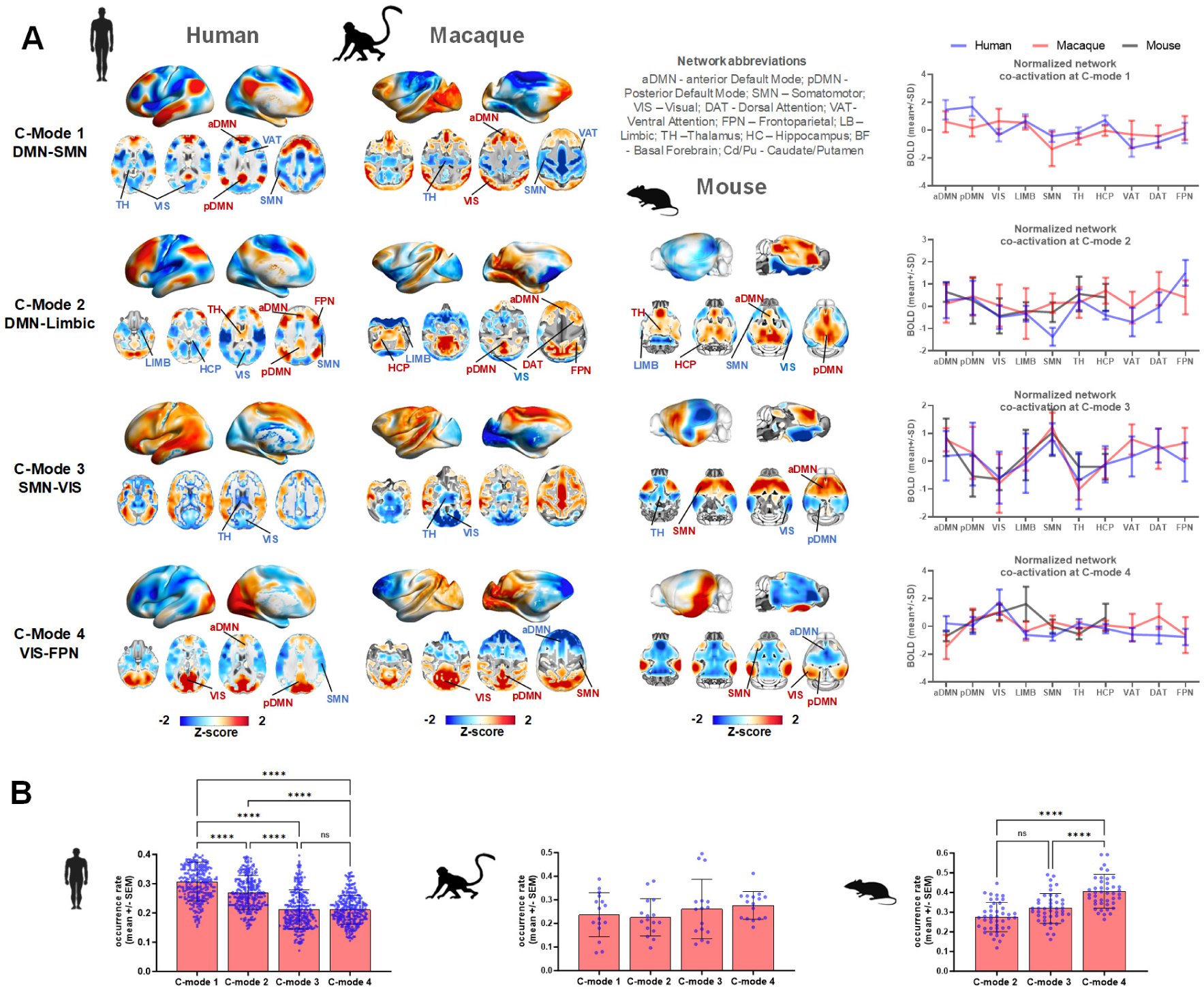
C-mode topography in awake humans, macaques and mice. **(A)** Z-scored C-mode maps (Left) and corresponding normalized network co-activation profiles (Right, mean +/- SD of voxels within the network mask). **(B)** C-mode occurrence rates (mean +/- SEM), and between C-mode comparisons (Kruskal-Wallis test, FDR corrected). Abbreviations: aDMN - anterior Default Mode; pDMN - Posterior Default Mode; SMN – Somatomotor; VIS – Visual; DAT - Dorsal Attention; VAT-Ventral Attention; FPN – Frontoparietal; LIMB – Limbic; TH –Thalamus; HC – Hippocampus; BF - Basal Forebrain; Cd/Pu - Caudate/Putamen. P-values: *p < 0.05, **p < 0.01, *** p < 0.001, **** p < 0.0001.

Collectively, cross-species C-mode matching revealed 4 topographically related C-modes in human and macaque, three of which (C-modes 2-4) were also represented in the rodent brain (**Figure 2A**). In all species, C-modes exhibited rich spatial organization encompassing positive and negative peaks of BOLD activity that together delineated a set of stereotypic network coactivation profiles (**Figure 2A**, right panels). Specifically, C-mode 1 *(DMN-SMN)* encompassed peaks of BOLD activity in DMN, accompanied by below baseline activity in SMN and ventral attention VAT areas in both humans and macaques. C-mode 2 (*DMN-Limb*ic) exhibited coactivation of DMN areas in anti-correlation with SMN, VIS, and LIMB networks in all three species. Interestingly, while in humans the HCP had below-baseline fMRI activity, macaques and mice show above-baseline fMRI activity in this region. C-mode 3 (*SMN-VIS*) activity peaked in the SMN and concomitantly engaged most cortical regions of the human brain, albeit with considerably weaker or negative BOLD activity in VIS, TH and basal forebrain areas, reminiscent of global fMRI signal (GS) fluctuations. Finally, C-mode 4 (*VIS*-*FPN*) was characterized in humans and macaques by positive coactivation in VIS and posterior cortical regions, and negative coactivation in FPN cortical regions. This topography in mice and macaques, but not humans, was associated with co-deactivation of anterior cingulate and prefrontal regions of the DMN. Taken together, these results point at the presence of notable topographic correspondences in the functional organization of dominant C-modes in human, macaque and mouse brains.

We next computed for each C-mode its occurrence rate, defined here as the proportion of fMRI frames assigned to each C-mode for each subject/animal. Interestingly, while the spatial organization of C-modes exhibited species-invariant topographic features, their occurrence rate, showed variation across species (**Figure 2B**). Specifically, in humans we observed a dominant occurrence of DMN-SMN C-mode 1 and 2 (Kruskal-Wallis test, p < 0.001) while mice occurrence of sensory-oriented C-mode 3 and 4 was observed instead (Kruskal-Wallis test, p < 0.001), with macaques showing equiprobable (Kruskal-Wallis test, p = 0.25) C-mode occurrence (although a slight trend for a mouse-like profile was apparent). Thus, the temporal structure of C-modes in humans was biased towards a greater occurrence of polymodal integrative metastates (C-modes 1 and 2), with mice (and possibly macaques= exhibiting instead a greater occurrence of sensory-oriented network modes (C-modes 3 and 4).

### fMRI C-modes exhibit infraslow fluctuations in humans, macaques and mice

Our formalism allowed us to compare C-Modes with single fMRI frames and thus uncover the temporal structure of the spatial configurations that underlie spontaneous fMRI dynamics. We leveraged this property to describe, in all species, the temporal evolution of each C-mode for each subject/animal by computing the instantaneous spatial correlation between each C-mode map and each fMRI frame in all timeseries. The power spectra of the resulting “C-mode timeseries” revealed that C-modes undergo infraslow fluctuations in all species, with most of the power peaking within the 0.01-0.03 Hz range (**Figure 3**). Peaks of infraslow activity were distinct and sharp in humans, and slightly less prominent (yet clearly recognizable) in mice and macaques. We next investigated the assembly of each C-mode by normalizing (z-score) its time-course and by time locking peaks of C-mode time courses (i.e. event-wavelet) at the group level. We found that C-modes in awake humans, macaques and mice assemble and disassemble in a slow and gradual fashion (**Figure 3, red insets**), reminiscent of damped oscillations in the dominant frequency band. These results suggest that fMRI C-modes exhibit comparable infraslow dynamic cycling in all the examined mammalian species.

**Figure 3.**
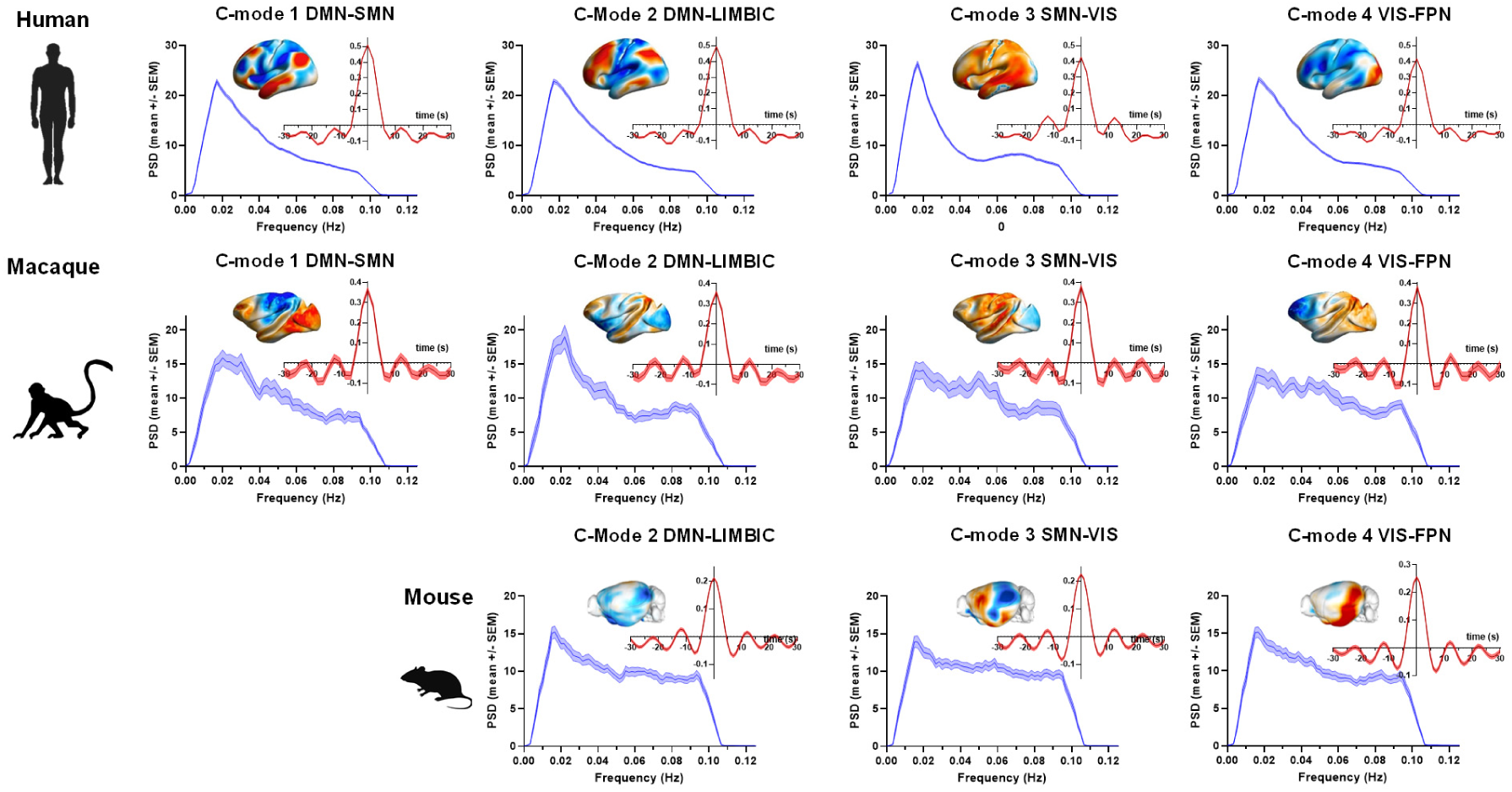
Infraslow dynamics and formation of C-Modes. Group-level power spectral density (blue, mean+/- SEM) of C-Mode to fMRI frame correlation time-series. Red insets denote the mean+/- SEM correlation values time-locked to peaks in the C- Mode time-series.

**Figure 4.**
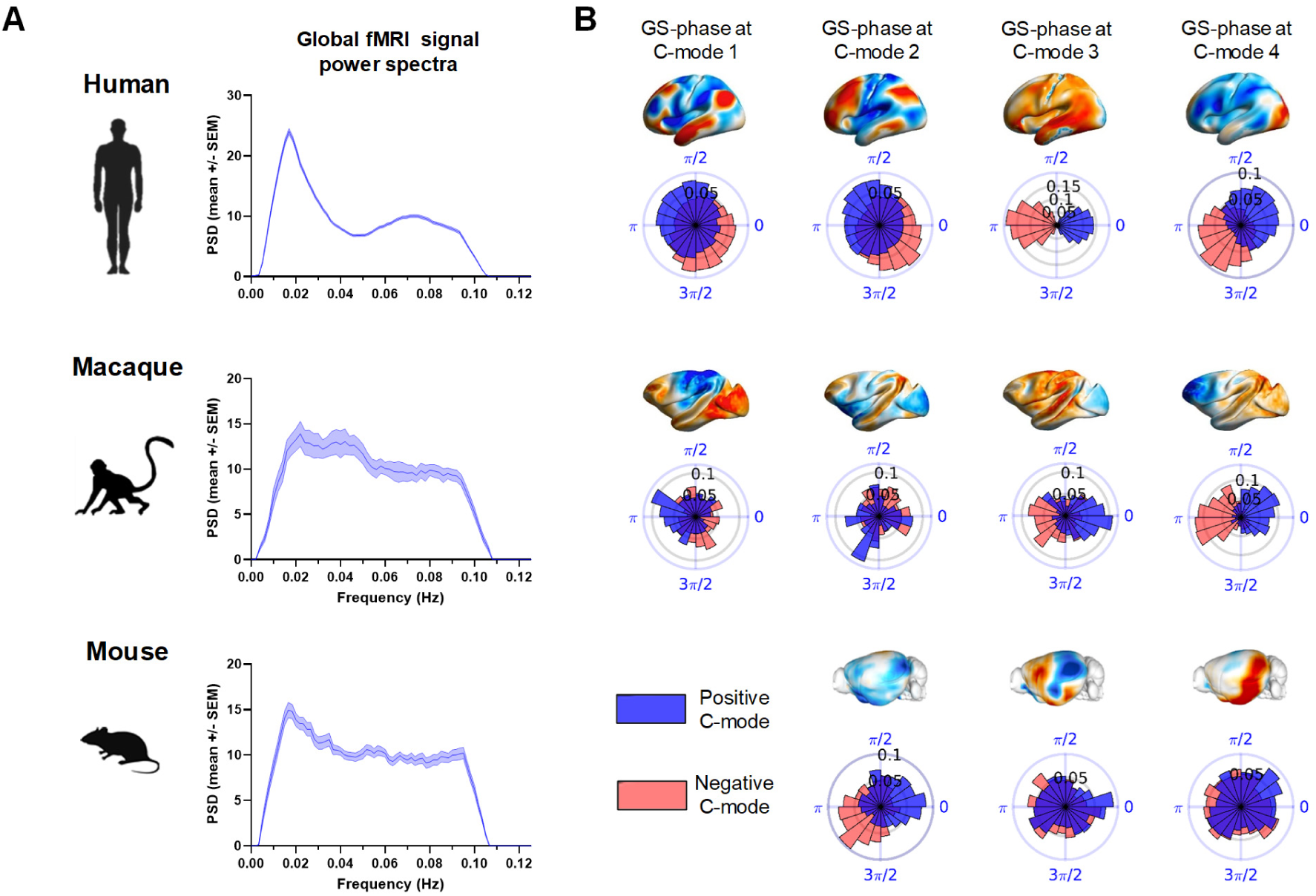
C-mode occurrence within fMRI global signal infraslow cycles. **(A)** Group-level power spectral density (mean+/- SEM) of the GS. **(B)** Distribution of GS-phases at the occurrence of each C-mode. Blue and red distributions correspond to GS phases sampled from the positive and negative C-mode time courses, respectively. All distributions significantly deviate from circular uniformity (Rayleigh test, p < 0.05, FDR corrected).

### fMRI C-modes occur at specific phase of fMRI global signals cycles in humans, macaques and mice

Prompted by previous investigations in anesthetized mice^16,30^, we next probed if C-modes in higher species have a preferred occurrence within fMRI global signal (GS) cycles. We thus first ascertained that also in awake conditions the GS would dominantly fluctuate within the infraslow range in all species (**Figure 4A**). A spectral analysis showed that the power spectrum of GS sharply peaks within a 0.01-0.03 Hz band in awake humans, with analogous (albeit less pronounced) peaks of activity in the same frequency range in both awake macaques and mice. We then built a circular distribution of the phases of the filtered (0.01-0.03 Hz) GS at which each C-mode occurred, by sampling only occurrences in which the normalized C-mode time series surpassed threshold values of 1 SD. C-mode occurrence was significantly phase-locked within GS cycles in humans, macaques and mice (**Figure 4B**, Rayleigh test, p < 0.05, FDR corrected). Interestingly, GS-phase distributions in macaques and humans presented key similarities, with C-modes 1, 3 and 4 (but not 2) exhibiting remarkably conserved cross-species phase alignment. Moreover, C-modes 3 and 4 showed broadly similar circular means across all the three species examined here.

C-modes were computed by coalescing a CAP with an anti-CAP of nearly identical spatial shape but opposite polarity. This description of brain dynamics works well under the assumption that CAP and antiCAPS represent a single fluctuating brain sub-state whose pattern of activity changes sign cyclically. To corroborate the cyclic nature of C-mode fluctuations within GS cycles, we repeated the sampling of GS phases but with inverted (negative) C-mode timecourses so to capture the full temporal evolution of C-mode. This approach revealed clearly opposite distributions of sampling of the GS phases from positive and negative C-mode occurrences (**Figure 4B, red insets**). These results suggest that C-modes describe cycling spatiotemporal sub-state that fluctuates in magnitude and sign according to the infraslow structure of the global fMRI signal.

To further provide evidence of the infraslow oscillatory nature of C-mode dynamics, we investigated whether different C-modes were phase-coupled within individual GS cycles. We computed the GS phase difference between the occurrences of a C-mode in a GS-cycle, and following occurrences of another C-mode either within the same GS-cycle, or in an immediately subsequent cycle (**Figure S7**). These analyses showed highly consistent positioning of a C-mode within each GS cycle (**Figure S7**, diagonals) and phase coupling between some C-mode pairs (see C-mode 2 and 3 and 3 and 4 in humans, and C-modes 3 and 4 in mice). These results suggest that in mammalian species, intrinsic fMRI signal fluctuations do not reflect spatially undifferentiated peaks and troughs of BOLD activity, but instead encompass infralow cycling between dominant patterns of BOLD activity. This general principle can be extended to entail the evolutionary conservation of the phase-relationship with fMRI GS cycles for most (albeit not all, see C-mode 2) of the explored C-modes. In sum, these results show that C-modes are phase locked to intrinsic GS fluctuations and imply that different C-modes can be conceptualized as networks of coupled oscillators in multiple mammalians species.

### Coupled oscillatory activity explains C-modes transition dynamics

We next considered whether the temporal structure of C-mode instantaneous transitions could similarly be underpinned by species-invariant principles. For each species, we modeled the system as a Markov process from sequences of concatenated C-mode occurrences and computed the transition probability into a different C-mode, as well as C-mode self-transitions (also termed persistence probability, **Figure 5A**). We found that the most recurring C-modes (1-2 in humans, 3-4 in macaques, and 3-4 in mice) were sinks of preferred directional transitions (p < 0.01, black crosses in **Figure 5A**), which can as such considered as state attractors.

**Figure 5.**
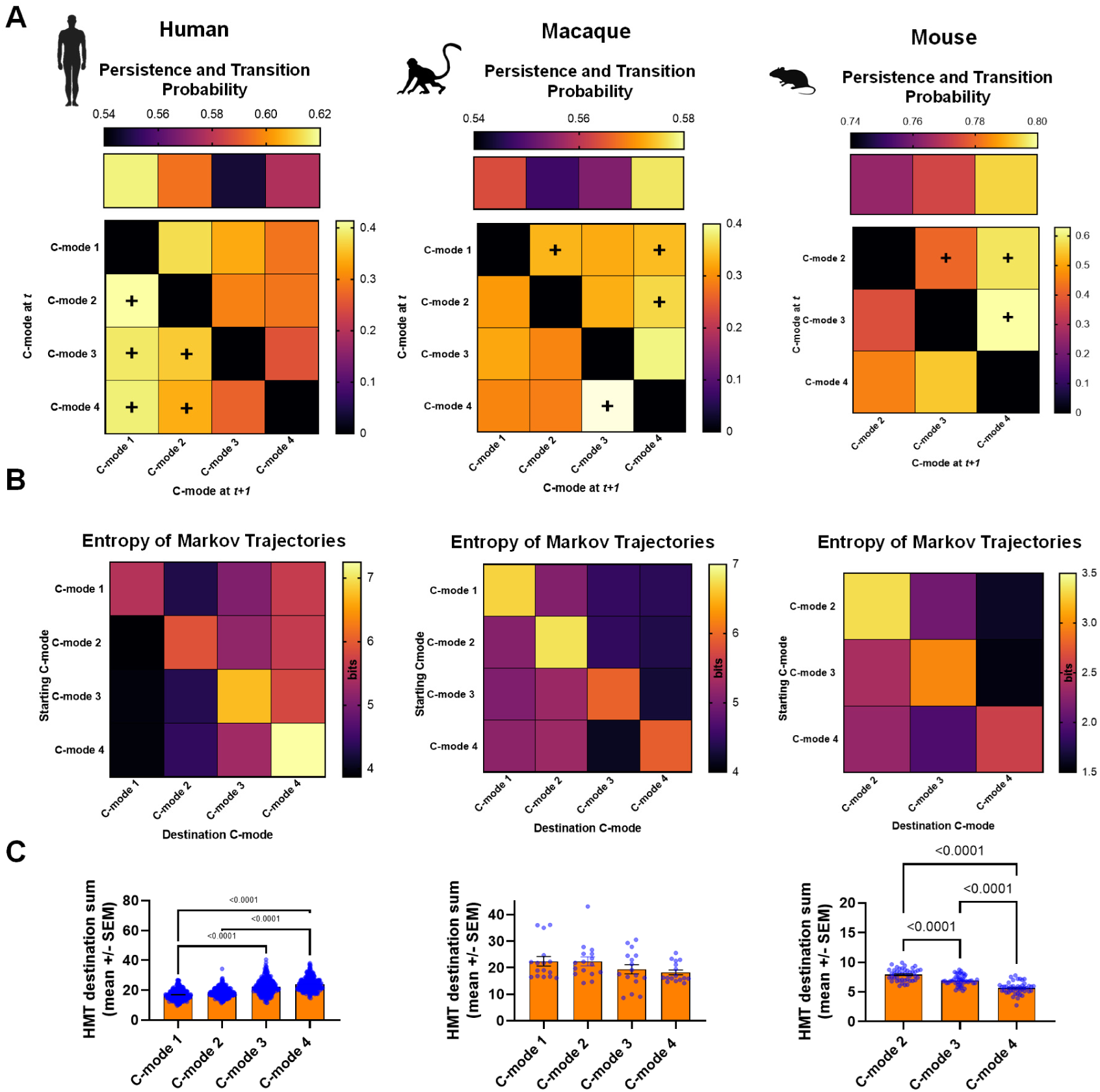
Temporal trajectories of C-modes converge to the most recurring state. **(A)** Persistence (top row) and transition (off- <u>diagonal) probability in human (left), macaques (middle), and mice (right). Red crosses denote the destination C-mode of</u> <u>preferred</u> directional transitions (Pij > Pji). **(B)** Entropy of Markov trajectories (HMT) show that the C-mode with the highest accessibility are C-modes with higher occurrence rates (i.e. C-mode 1 in humans, and C-mode 4 in macaques and mice). Higher entropy indicates lower accessibility of a destination C-mode (column) from a starting C-mode(row). **(C)** Quantification of the sum of Entropy of Markov Trajectories for all destinations (columns in the HMT matrices at the single-subject level), and comparison between the means (one-way ANOVA, and Tukey test for multiple comparisons). The most occurring C-modes (cf **Figure 2**) are also the most accessible ones. P-values: *p < 0.05, **p < 0.01, *** p < 0.001, **** p < 0.0001.

Furthermore, we investigated, for each species, the “accessibility” of C-modes from one other by computing the corresponding Entropy of Markov Trajectories (HMT)^33^ from the transition probability matrices. This parameter measures the complexity of a transition: low entropy values imply an almost deterministic direct path or high accessibility. On the contrary, high entropy values suggest high uncertainty, requiring random steps through different C-modes before reaching the destination, i.e. low accessibility. In keeping with the C-mode occurrence rates we describe in **Figure 2B**, the most recurring C-modes in all species were also those that were most accessible, i.e they were the one characterized by lowest entropy values (**Figure 5B**, C). Specifically, in humans, C-modes 1 and 2 were the most accessible ones (p < 0.0001 against C-modes 3-4). In macaques we did not find any C-mode to have preferred accessibility (p = 0.25) Conversely, in mice, C-modes 3 and specially 4 were the most accessible ones (p < 0.0001). Importantly, in all species the ensuing accessibility profile (**Figure 5C**) recapitulated the C-mode occurrence rates we described in **Figure 2B**, with the most accessible C-modes being also the most occurring ones (**Figure 2B**). This was true also in macaques, where the entropy of Markov trajectories required to reach C-modes 3 and 4 followed the tendency of higher occurrence rates of these C-modes.

In the sections above we characterized intrinsic fMRI dynamics both in terms of coupled quasi-periodic fluctuations between coactive networks captured by different C-modes, and also in terms of transitions between C-Modes. If coupled cyclic activity is key to explaining state transition dynamics, we expect that trajectories between C-modes with lower entropy (i.e., small HMT, corresponding to more direct transitions) would occur on average with shorter infraslow phase differences. Conversely, C-mode pairs with larger entropy (i.e. higher HMT corresponding to less direct transitions) would occur on average with longer infraslow phase differences. To test this hypothesis, we computed the circular-linear correlation between the mean GS-phase differences between C-mode occurrences (**Figure S7**) and the HMT for their trajectories. We found clear and significant correlation values in all species (r = 0.66, 0.66, 0.87 with p = 0.029, 0.031 and 0.032 for humans, macaques and mice respectively). This implies that the transition structure of C-modes is in part described by infraslow coupled cyclic dynamics across all species, embedding fast transition phenomena within GS cycles in the dominant infraslow band.

### fMRI C-mode occurrence predicts ranking of connectivity gradients in humans, macaques and mice

Previous investigations have shown that a high portion of the variance in static fMRI connectivity is explained by a limited fraction (5-15%) of fMRI frames exhibiting exceedingly high cofluctuation amplitude^16,34,35^. We thus investigated whether the dynamic occurrence of dominant C-modes alone could similarly be sufficient to reconstitute key organizational features of the static fMRI connectome. To this aim, we first examined whether the occurrence-weighted average of C-modes would reproduce the static architecture of fMRI connectivity. To this purpose, we calculated for each C-modes its co-fluctuation matrix, i.e. the cross-multiplication of each C-mode map with itself^36^. We found that in all species, the weighted average by occurrence rate of C-mode co-fluctuation matrices exhibited high correlation with the corresponding group-mean static fMRI connectivity matrix (r > 0.57, all species, with r = 0.75 in humans, **Figure S8**). These findings corroborate the notion that C-modes dynamics account for high co-fluctuation events critical for the topographic organization of the static functional connectome.

To further investigate the relationship between C-mode dynamics and static fMRI connectivity, we inquired whether C-mode occurrence could also be related to the organization of fMRI connectivity gradients^37,38^. Here we posited that C-mode occurrence rate may be linked to the ranking of functional connectivity gradients, with dominant (i.e. most occurring) C-modes aligning with the gradients that explain the most variation in fMRI connectivity. To test this hypothesis, we first computed for each species the top five fMRI connectivity gradients, which we next ranked by decreasing variance explained (**Figure S9**). We then compared, for each species, the spatial similarity of the obtained gradients to that of each C-mode, matching them according to their highest absolute spatial correlation. Supporting our hypothesis, plausible spatial correspondences between C-mode and gradient topographies were observed in all species (**Figure 6A**). Specifically, in humans the most occurring C-modes 1 and 2 were matched with dominant gradients 1 and 2, while in macaques and mice dominant gradients 1 and 2 were matched with most occurring C-modes 4 and 3. Moreover, in all species C-mode occurrence was linearly related to the variance explained by each gradient (R^2^ > 0.78 for all species, **Figure 6B**). This analysis shows how recurring network interaction represented by dominant C-modes, and their relative occurrence rate, shape the organization of the static fMRI connectome and its principal axis of variance in all the probed species.

**Figure 6.**
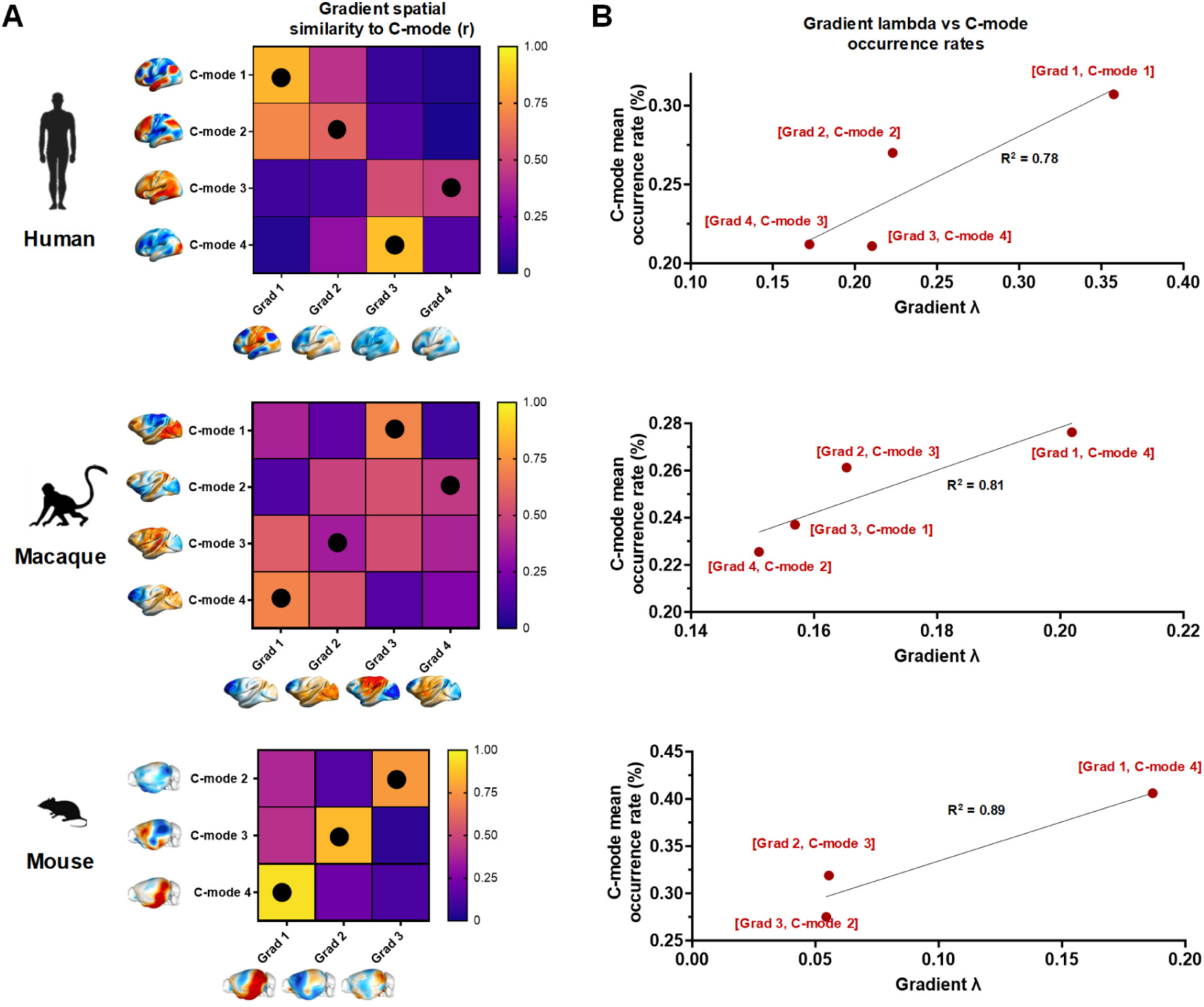
C-modes occurrence rate predicts ranking of functional connectivity gradients. **(A)** Spatial correlation between the principal gradients and each C-mode map. Black dots denote the spatial matching between maps obtained using the Hungarian Algorithm. **(B)** Scatter plot of the variance explained by each gradient (lambda) versus the corresponding C-mode occurrence rate. R-square from a linear fit (p < 0.01). The most occurring C-modes accounts for most variance in the corresponding gradient axis.

## DISCUSSION

While previous cross-species studies have attempted to compare the static functional architecture of specific fMRI networks via time-invariant fMRI connectivity mapping^8,39,40^, whether the dynamic organization of intrinsic fMRI activity is evolutionarily conserved remains unclear. To fill this knowledge gap, we performed a first-of-its-kind systematic investigation of intrinsic fMRI dynamics in awake humans, macaques and mice using the same analytical framework. To facilitate cross-species comparisons, we introduced a parsimonious description of network dynamics via fMRI C-modes, which represent dominant patterns of fluctuating BOLD activity. Using this simple approach, we found that fMRI dynamics in awake humans, macaques and mice encompasses the recurring occurrence of a set of functionally-related C-mode topographies. We also report that the dynamic structure of fMRI network activity follows a set of evolutionarily-invariant principles. These include the observation that C-modes undergo infraslow fluctuations and tend to occur at specific phases of the fMRI global signal. Moreover, their transition structure is partly explained by infraslow coupled oscillator dynamics within fMRI global signal cycles. We also show that C-mode occurrence accounts for high co-fluctuation events critical to the topographic organization of the static functional connectome, and is closely related to the ranking of connectivity gradients. These observations suggest that, beyond and above some expected species-specific features, the dynamic structure of intrinsic fMRI activity in the awake mammalian brain follows evolutionarily conserved principles.

Although the lack of systematic comparisons of the dynamic organization of fMRI activity across species does not allow us to directly relate our findings to prior literature, our results are consistent with emerging evidence supporting the presence of cross species homologies in static organization of fMRI connectivity in the mammalian brain^31,41^. Our results expand these initial investigations, by showing that correspondences in static fMRI network organization can be parsimoniously described and accounted for by a limited set of evolutionarily-related dynamic patterns of BOLD activity. In keeping with this, the topographic organization of C-modes encompasses peaks of BOLD activity spanning network systems previously described in multiple mammalian species, including components of the DMN, salience and motor-sensory networks, as well as in many key subcortical systems^10,28,30^. Extending previous observations^30,42,43^, we also found that C-modes dynamics can be reliably related to (and as such, it “explains”) the static organization of the fMRI connectome in all the probed species. These finding reconcile prior investigations of the dynamic structure of fMRI connectivity in rodents^16,44–46^, macaques^47^, and humans^15,17,27,48–52^ by showing that C-mode dynamics encompass high-amplitude peaks of BOLD activity that critically shape the steady-state architecture of the fMRI connectome. These results are also in agreement with the assumption that the mechanisms underlying interareal information transfer as assessed with fMRI are conserved in the mammalian brain^53,54^.

Our observation that the dynamic structure of fMRI activity in mammals follows species-invariant principles is important in the light of the notion that many fundamental physiological and anatomical features (including the brain’s anatomical architecture), are known to scale with body size, or to be marginally (or not) represented in lower species^55–57^. Our data suggests instead that, independent of evolutionary complexity, infraslow fluctuations of spatially rich patterns of fMRI activity similarly underpin spontaneous brain activity in multiple mammalian species. In this respect, one important advancement of the present study is the use of datasets collected in awake conditions in all species, an experimental strategy that allowed us to carry out a cross-species comparison of fMRI dynamics unconfounded by the pharmacological effects (and the ensuing brain state changes) produced by anesthesia. An additional key benefit of the C-mode framework we introduce here is computational tractability, which allows for the representation of the corresponding spatiotemporal patterns of fMRI activity with voxel resolution (i.e. without predefined anatomical boundaries) and avoiding the use of correlation-based metrics (e.g. like in sliding window-based dynamic connectivity mapping). Moreover, the employed approach allows for a fine-grained temporal localization of dynamic effects with single-frame resolution. All of these properties were key to the identification of the dynamic properties and cross-species correspondences we report in this work.

The voxelwise topography of C-modes enabled us to match and compare these spatiotemporal patterns at maximum spatial resolution, revealing a set of anatomically related motifs that exhibit evidence of evolutionary conservation across species. Functional correspondences between human and macaque were apparent, and encompassed four neuroanatomically homologous C-modes that were plausibly identified in two independent human datasets. While evolutionarily more tentative, spatial matching between macaque and mouse was also plausible, with preserved fMRI polarity in key anchor networks in the three matched C-modes, and evidence of conserved C-mode GS phase coupling in both species. Improved anatomical matching may be obtained in future studies by incorporating data from lissenchephalic new world monkeys, and other primate species phylogenetically closer to rodents that could serve as intermediate evolutionary link between macaques and mice^6,58,59^. This approach may represents a warranted extension of our work, owing to the increased availability of fMRI data in multiple primate species^6^.

The observation of a substantial coupling between C-mode occurrence and fMRI global signal cycling was first reported in anesthetized mice^16^ and it is here extended to awake humans and macaques. This finding corroborates the hypothesis that fluctuations in fMRI GS are the intrinsic manifestation of highly structured network interactions^60^. This results is also of interest in the light of emerging evidence linking global fMRI activity to intrinsic fluctuations in arousal^27,61–63^. Within this framework our findings suggests that the cycling spatiotemporal patterns of activity captured by C-modes (or by other analogous computational frameworks, like quasi periodic patterns^27,34^), could be strongly affected or driven by ascending modulatory transmission.

Over the last decade, several influential mathematical models have described resting state activity in terms of networks of coupled oscillators^21^. This work provides important empirical support to this modelling as it shows, from data and without making assumptions about the mechanism, that transition probabilities between different brain states can be described in terms of coupled C-Modes oscillating at infraslow frequencies. Moreover, the fact that the state transitions can be reliably described as coupled oscillators in all species suggests that a functional architecture based on coupled oscillating networks offers important evolutionarily advantageous computational benefits. These include the possibility of rapidly reconfiguring coordination and communication between different brain regions, and effectively transferring information across scales and brain areas^21,54,64^.

While the present work focuses on the description of species-invariant principles underling the organization of spontaneous fMRI activity, species-specific features were also apparent. Beyond foreseeable discrepancies in the topography of specific C-modes, which can be related to the increased complexity of the cortical mantle across the phylogenetic tree, one interesting difference we observed is a shift in C-mode occurrence across species. Although comparable human C-modes were identified in macaques and mice, their occurrence rate was inversed, with C-modes 1-2 being the most occurring patterns in humans and C-modes 3-4 in mice and in macaques. Taken together, these observations support the evolutionary basis for ongoing dynamic changes across species, albeit with a possible shift in the time that each species spends in each spatiotemporal state. Such species-dependent shift in C-mode occurrence may reflect brain adaptations that support the increasing demands of higher-order cognition throughout evolution. The finding that in humans, BOLD activity profiles in C-modes 1-2 and 3-4 peak in polymodal and sensory areas, respectively, suggests that the intrinsic organization of fMRI activity is biased towards introspective states that are less frequently visited in animals. Accordingly, we found that these most recurrent C-modes are also the most easily accessible from any other C-mode. Our results also highlight a link between the dynamic organization of fMRI activity and the principal axis of variance of static fMRI activity as mapped with functional connectivity gradients. This result suggests that the macroscale organization of the functional connectome is critically shaped by the occurrence of its constituting spatiotemporal modes, a notion supported also by complementary conceptualization of fMRI dynamics^65^.

In summary, we describe a conserved set of dynamic rules governing large-scale fMRI network dynamics in human, macaques and mice. Our work provides a simple and flexible framework to quantitatively model and relate intrinsic fMRI dynamics across the phylogenetic tree.

## MATERIALS AND METHODS

### Data and preprocessing

Resting state fMRI (rsfMRI) datasets from awake, freely breathing humans, macaques, and mice were used in this study. Preprocessing included most steps suggested by the guidelines of the Human Connectome Project^66^, using a combination of fMRI dedicated software AFNI^67^, FSL^68^, FreeSurfer^69^, and SPM12 (http://fil.ion.ucl.ac.uk/spm/).

#### Human

The main dataset, Hangzhou Normal University of the Consortium for Reliability and Reproducibility (CoRR-HNU, or HNU)^23^ includes 10 sessions of 10-min scans over the course of a month from n = 30 young healthy adults with no history of neurological or psychiatric disorders, head injuries, nor substance abuse (balanced sexes, age = 24+/- 2.41 years). Before scanning, participants were asked to relax and remain still with their eyes opened, avoiding falling asleep. During scanning, a black crosshair was shown in the middle of a grey background. The study was approved by the ethics committee of the Center for Cognition and Brain Disorders at Hangzhou Normal University, and all participants signed written consent before data collection. A GE MR750 3T scanner (GE Medical Systems, Waukesha, WI, USA) was used to acquire MRI data. Functional scans were acquired with an echo-planar imaging sequence - EPI: TR = 2 s, TE = 30 ms, flip angle = 90°, FOV = 220 × 220 mm, matrix = 64 × 64, voxel-size = 3.4 mm isotropic, 43 slices. Data was downloaded from the International Neuroimaging Data-Sharing Initiative (INDI - http://fcon_1000.projects.nitrc.org/indi/CoRR/).

The Midnight Scan Club (MSC) dataset^24^ was used as secondary, replication dataset. Five out of 10 randomly selected sessions were used from separate days, and included 30-min scans from n = 10 healthy young adults (balanced sexes, age = 29.1 +/- 3.3 years). Participants were asked to visually fix on a white crosshair against a black background. The study was approved by the Washington University School of Medicine Human Studies Committee and Institutional review Board, and all participants signed written consent before scanning. Functional scans were acquired with a Siemens TRIO 3T MRI scanned (Erlangen, Germany) using a gradient-echo EPI sequence: TR = 2.2 s, TE = 27 ms, flip angle = 90°, voxel-size = 4 mm isotropic, 36 slices. Data was downloaded from OpenNeuro (doi:10.18112/openneuro.ds000224.v1.0.3).

Preprocessing. The first 5 fMRI volumes were removed from each subject’s raw data, then despiking (AFNI *3dDespike*) and slice-timing correction (AFNI *3dTshift*) was performed. Data subsequently underwent motion-correction (AFNI *3dvolreg*); skull-stripping (FSL *fast* and *bet*^70^); co-registration to the MNI 3 mm isotropic template (FSL *flirt*); regression of nuisance parameters (white matter, cerebrospinal fluid, and 24 motion parameters (6 parameters, 6 derivatives, and their respective squared time-series) (AFNI *3dDeconvolve*); band-pass filtering between 0.01-0.1 Hz (AFNI *3dBandpass*); spatial smoothing with a 6 mm FWHM kernel (AFNI *3dBlurInMask*); and voxel time-series were finally normalized to z-scores (zero-mean, and standard deviation units).

#### Macaque

The Newcastle dataset (NC) includes n = 14 rhesus macaque monkeys (*Macaca Mulatta*) scanned with no contrast agents, from which N = 10 animals (2 females, 8 males), in which two independent fMRI session were available, were used for our analyses (age = 2.28 +/- 2.33, weight = 11.76 +/- 3.38). Two additional (female) animals were discarded from our analyses due to excessive head motion exceeding, in either session, over 30% of fMRI volumes with Framewise Displacement above a 0.3 mm threshold. Animal procedures, head-fixation, and protocols were approved by the UK Home Office and comply with the Animal Scientific Procedures Act (1986) on the care and use of animals in research and with the European Directive on the protection of animals used in research (2010/63/EU) (see Slater et al., 2016 for protocol specifics on animal preparation for awake imaging^25^). A Vertical Bruker 4.7T primate dedicated scanner was used and rsfMRI experiments were performed in awake, head-fixed animals for two separate sessions with TR = 2 s; TE = 16 ms, voxel-size = 1.2 mm isotropic. Data was downloaded from NHP data sharing consortium PRIME-DE (http://fcon_1000.projects.nitrc.org/indi/indiPRIME.html)^26^.

Preprocessing. The first 5 rsfMRI volumes were removed from each animal’s raw data, then despiking (AFNI *3dDespike*) and slice-timing correction (AFNI *3dTshift*) was performed. Data subsequently underwent motion-correction (AFNI *3dvolreg*); skull-stripping (FSL *fast* and *bet*^70^); co-registration to the Yerkes19 2 mm isotropic template^71^ (FSL *flirt*); regression of nuisance parameters (white matter, cerebrospinal fluid, and 24 motion parameters (6 parameters, 6 derivatives, and their respective squared time-series) (AFNI *3dDeconvolve*); band-pass filtering between 0.01-0.1 Hz (AFNI *3dBandpass*); spatial smoothing with a 3 mm FWHM kernel (AFNI *3dBlurInMask*); and voxel time-series were finally normalized to z-scores (zero-mean, and standard deviation units).

#### Mouse

C57BL/6J mouse data was obtained from n = 44, head-fixed awake male mice undergoing a 12-min rsfMRI scan using the same animal preparation, habituation and scanning protocols previously described^30^. *In vivo* experiments were conducted in accordance with the Italian law (DL 26/214, EU 63/2010, Ministero della Sanita, Roma) and with the National Institute of Health recommendations for the care and use of laboratory animals. The animal research protocols for this study were reviewed and approved by the Italian Ministry of Health and the animal care committee of Istituto Italiano di Tecnologia (IIT). All surgeries were performed under anesthesia. Young adult (< 12 months old) male C57BL/6J mice were used. RsfMRI scans, both retrieved and newly acquired, were acquired at the IIT laboratory in Rovereto (Italy) using a Bruker 7T scanner (Bruker Biospin, Ettlingen) with a BGA-9 gradient set, 72 mm birdcage transmit coil, and a four-channel solenoid receiver coil: TR = 1 s, TE = 15 ms, flip angle = 60°, matrix = 100 × 100, FOV = 2.3 × 2.3 cm, 18 coronal slices 0.6 mm thick, 12 minutes total acquisition time.

Preprocessing. The first 120 rsfMRI volumes (2-min) were removed from each animal’s raw data to account for thermal gradient equilibration, then despiking (AFNI *3dDespike*) was performed. Due to the short TR we did not perform slice-timing correction in these images. Data subsequently underwent motion-correction (FSL *mcflirt*); skull-stripping (FSL *fast* and *bet*^70^); co-registration to an in-house mouse brain template of 0.23 × 0.23 × 0.6 mm^3^ (ANTS registration suite^72^); regression of nuisance parameters (white matter, cerebrospinal fluid, and 24 motion parameters (6 parameters, 6 derivatives, and their respective squared time-series) (AFNI *3dDeconvolve*); band-pass filtering between 0.01-0.1 Hz (AFNI *3dBandpass*); spatial smoothing with a 0.5 mm FWHM kernel (AFNI *3dBlurInMask*); and voxel time-series were finally normalized to z-scores (zero-mean, and standard deviation units).

### Whole brain CAP detection and cluster-number selection

To identify recurrent rsfMRI whole-brain states, we used the whole-brain coactivation patterns (CAPs) approach^15–17^ in which fMRI frames are cluster based on their spatial similarity and then averaged to define recurrent patterns of BOLD coactivation. Specifically, for each species, we first performed censoring of motion-contaminated frames (framewise Displacement: FD > 0.3, 0.3, and 0.075 mm for humans^2^, macaques^29^, and mice^30^ respectively), and then concatenated the frames from all subjects or animals. Given that clustering human fMRI data in the 3 mm MNI-space became computationally challenging owing to its large dimensionality (n-voxels = 43.539, compared to 11.402 in macaques, and 8.937 in mice), this step was carried out upon reducing data using the coarse 950-ROI Craddock Parcellation^73^. The choice of this specific parcellation regards its ability to provide a fair dimensionality reduction, while preserving information present at the voxel scale^73^. After these final steps, we ran, for each species, the k-means clustering algorithm^16,74^ (spatial correlation as distance metric, 500 iterations, 5 replications with different random initializations, from k = 2:20, 5 independent runs). CAP maps were obtained at the group level by averaging the fMRI frames belonging to a cluster at the voxel level, then normalizing these values to T-scores from the concatenated datasets (**Figure S1**). At the single-subject level, we obtained, for each subject/animal, a CAP-map for each cluster by only averaging and converting to T-scores, the frames belonging to a cluster but only within a subject/animal’s data/session. We note that CAP mapping through frame averaging in humans was done at the voxel level, as the parcellated data was only used for clustering purposes. After clustering, we recovered the censored fMRI frames to the cropped datasets and assigned them to the CAP with the highest spatial correlation. This was done in order to have a continuum of frames for subsequent analyses.

Selection of the optimal number of clusters was done following a set of previously proposed empirical rules^16,19^, as well as new metrics. These were dependent on the availability on test-retest sessions within a dataset in macaques and humans, as well as a full independent dataset in humans from a different site. Specifically, for human HNU and MSC datasets independently, we first ran the k-means clustering algorithm with the concatenated dataset from k = 2:20, selecting, for each ‘k’, the solution with the highest variance explained^16^ from 5 replications in 5 independent runs. Here, the variance of the data explained by each partition is defined as defined as the ratio between the between-cluster variance and the total variance (within-cluster + between-cluster variance). Within-cluster variance was computed as the averaged (over clusters) sum of square distances between elements in a cluster and its centroid. Between-cluster variance was computed as the averaged square distance between a cluster centroid and the centroid of all clusters or centroid of all data^16,75^. For each dataset independently, we first computed the variance explained by data partitioned into an increasing number of clusters (2 ≤ k ≤ 20) (**Figure S2A-B**). We next assessed the topographic consistency of CAPs at increasing partitions. To this purpose, we computed how consistent CAPs are by assessing, for each ‘k’ partition, the spatial correlation between a mean CAP map, and its matched map in the previous order (k-1) partition. Matching was done using the Hungarian Algorithm^32^. We found that partitions between k = 6:10 yielded, in both datasets, topographically stable CAPs that could be reliably identified at higher ‘k’ (**Figure S2C-D**).

We then assessed the within-dataset repeatability by comparing the spatial correlations between the mean CAP maps of each subject between each independent fMRI sessions (10 for HNU, and 5 for MSC, **Figure S2E-F**). Statistical significance of the mean within subject repeatability of each CAP was assessed by recomputing the spatial correlations between subject-level CAP maps after randomly shuffling the CAP-identity of fMRI frames, preserving occurrence rates. This process was repeated 1000 times, and repeatability values for each subject beneath the highest permutation value were flagged as non-repeatable (asterisks in **Figure S2E-F**). The result of these comparisons showed that within an upper limit of k = 13 (HNU) or 15 (MSC), all the mapped CAPs were represented in all fMRI sessions of each subject, with significant spatial correlation across sessions (p < 0.05, surrogate testing with randomly shuffled cluster associations). At higher partitions, one or more CAPs were instead no longer represented in one or more subjects. We further probed within-session stability of clustering by computing the CAP occurrence rate obtained across imaging sessions. We found that CAP occurrence rate (i.e. proportion of fMRI frames associated to a CAP in each subject), for both datasets, was stable across sessions at k = 2, 6, 8, and 9 (Kruskal-Wallis test, 10 groups for HNU, 5 groups for MSC, FDR corrected for k comparisons, **Figure S2 G-H**).

Finally, to maximize the generalizability of our partitioning, we compared the main dataset’s (HNU) CAP maps with those obtained in the MSC in terms of topography matching and occurrence rates. To this aim, we spatially compared mean group-level CAP maps (matched with the Hungarian Algorithm for each partition) and tested the significance of this comparison by recomputing the values after randomly shuffling the fMRI frames within each dataset, while preserving occurrence rates (**Figure S2I**). The mean occurrence rates from each dataset was assessed with a Wilcoxon signed rank test, p < 0.05, FDR corrected for k comparisons (**Figure S2J**). This analysis revealed that clusters from all partitions, except k = 6, 7, 9, 14, 15 and 16, were topographically reproducible across datasets (**Figure S2I**). By contrast, a comparison of the mean CAP occurrence rates between datasets (**Figure S2I**) showed that k = 8 was the only partition in which this parameter was conserved across dataset (Wilcoxon signed rank test, p < 0.05, FDR corrected for k comparisons). Based on these analyses, k= 8 was the only partition level meeting all the required within and between subject/dataset reproducibility criteria. We thus based all our subsequent analyses of human fMRI dynamics using k = 8 clusters.

Selection of optimal clusters in macaque fMRI time-series (Newcastle test-retest datasets, n = 8 animals, 2 sessions) followed the same strategy employed for human time-series. Briefly, computation of explained variance in macaque fMRI time-series revealed an elbow region within the k = 6:10 range (**Figure S3A**). Macaque CAPs were topographically stable at increasing k partitions until an upper limit at k = 10 (**Figure S3B**). Comparing repeatability of CAP topographies across sessions revealed that above k = 8, some CAPs present lack of topographical reproducibility (p > 0.05, surrogate testing with randomly shuffled cluster associations), as well as significant differences in CAP occurrence rates (Kruskal-Wallis test, FDR corrected for k comparisons, **Figure S3C-D**). This cumulative evidence suggests that k = 8 was the highest partition that guarantees CAP stability, as well as test-retest topographical and frame distribution repeatability amongst clusters.

For mice, the variance explained curve for the awake dataset we used in this work (N = 44) showed an elbow between 6 and 8 clusters (**Figure S3E-F**). Topographic CAP stability as a function of increasing partition number revealed that mouse CAPs were topographically stable up to k = 6. This value is in agreement with the results of CAP number selection in prior independent studies^30,40^, in which k = 6 was consistently identified as optimal partition in this species.

### Coactivation Modes and between species matching

In previous work we demonstrated that CAPs appear in pairs of spatial configurations with opposite BOLD coactivations^16^ (CAP, and anti-CAP, **Figure 1A**). As this feature was initially detected in awake and anesthetized mice^30^, we first showed that these anatomical configurations were present also in all of our datasets. For each species, we thus organized CAPs in descending occurrence rate and computed the Pearson’s correlation between the vectorized mean CAP maps as a spatial similarity metric, matching CAPs and anti-CAPs as pairs with the highest anticorrelation coefficient (**Figure 1B**). We then reduced pairs into C-modes, by taking each CAP map pair (T-score normalized), subtracting the least recurring one from its counterpart, then dividing the resulting map by two (**Figure 1A**). In order to have a normalized basis for between-species comparisons, we further z-scored each resulting C-mode map. Given that C-modes represent unique configurations between known resting state networks^16,17,19^, we extracted the mean BOLD values from voxels within masks of previously described, evolutionarily conserved^31,54^ cortical RSN in humans^28^, macaques^29^ and mice^30,76^, as well as hippocampal and thalamic masks^66,77^ (**Figure 1C**). For each C-mode, we computed their occurrence rates as the average occurrence of the CAP and anti-CAP conforming the C-mode. We then organized the vectorized network coactivation profiles of each C-mode in descending occurrence rate order, and first matched C-mode profiles of humans to macaques, then macaques to mice using the Hungarian Algorithm^32^ with vector-correlations between profiles as distance metrics for the cost function to be minimized (**Figure 1D**). We represented the similarities between network coactivation profiles of C-modes in the human HNU dataset and macaques (**Figure 1E**), as well as with the MSC dataset and macaques, confirming our matching (**Figure 1F**). Finally, we matched macaque C-mode network coactivation profiles with those of mice, without accounting for Fronto-Parietal (FPN), Ventral (VAT) and Dorsal Attention (DAT) networks, as these have not been described in the mouse brain (**Figure 1G**). We note that since C-mode 1 was not reliably identified in mice, most likely due to the lack of weighting by higher order VAT, DAT, and FPN, we hereafter exclude C-mode 1 from this species.

After matching and organizing C-modes, we mapped the voxel-wise C-modes as well as the network coactivation profiles within C-modes (within-network voxel mean +/- SD, **Figure 2A**). Independent group-level CAP maps for each human (HNU and MSC); macaque and mouse dataset are shown in **Figure S4**). For each subject/animal in each dataset, we computed a C-mode’s occurrence rate as the proportion of fMRI frames associated to the CAP pairs composing the C-mode. A comparison of the distributions of occurrence rates was then performed (Kruskal-Wallis test, FDR corrected).

We further tested that in-scanner head-motion did not selectively affect the configuration of any C-mode. This was done by first independently clustering the non-motion censored rsfMRI datasets with k = 8, 8, and 6 for humans, macaques and mice respectively; building C-modes; and then comparing the topographies of matched C-mode maps (Hungarian Algorithm^32^) to the ones obtained after censoring (Pearson’s Correlation) (**Figure S5A**). We then computed, for each C-mode in each subject/animal, the proportion of frames in it that were classified as high-head motion (FD > 0.3, 0.3, and 0.075 mm for humans, macaques, and mice, respectively), and compared these distributions across C-modes (one-way ANOVA, **Figure S5B**).

### C-mode infraslow dynamics, formation, and temporal structure

We investigated the temporal evolution of C-modes by generating a time-course of instantaneous C-mode to fMRI frame spatial correlation (hereafter referred to as “C-mode time-courses”) for each subject/animal, and then computing the power spectrum of these time-courses as well as that of the Global fMRI signal (GS, i.e., the average of all voxels at each frame). We next computed the group mean +/- SEM power spectra. C-mode time-courses were normalized to standard deviation units and mean zero. The evolution of the formation of a C-mode was computed by time-lock averaging the C-mode to frame correlation values at the vicinity of a peak surpassing 1SD (+/- 30 seconds), then averaging these event-kernels at the group level (**Figure 3 insets**).

Given that the GS also presented a strong peak in the power spectrum in the [0.01-0.03Hz] infraslow band (**Figure 4A**) we computed whether occurrence of C-modes was phase locked to infraslow GS fluctuations^16,30^. We extracted the instantaneous infraslow phase of the GS (filtered between 0.01-0.03Hz) using the Hilbert Transform, then divided the trace into cycles of minimum 30 s and maximum 100 s. For each cycle, if a C-mode was present, we sampled the GS-phase at which the C-mode occurred, retaining only samples from normalized C-mode time-courses exceeding 1SD, thus guaranteeing the selection of frames that are reasonably well assigned to a specific C-mode^16^. We then built the distribution of GS-phases from the group-concatenated sampling of C-mode occurrences, and using MATLAB’s CircStats toolbox^78^, we performed a Rayleigh test (p < 0.001, FDR corrected) for deviations from circular uniformity (**Figure 4B**). To confirm the cyclical features of C-mode fluctuations within GS cycles, we repeated this analysis but sampling GS phases from occurrences of inverted (negative) C-mode time courses (**Figure 4B, red insets**). As in previous our work^16^, we also demonstrated that C-modes are phase-locked within GS-cycles by sampling the GS-phase difference between the occurrence of C-modes within a GS cycle, and the following occurrences of another C-mode within the same cycle or the immediately subsequent one, limiting sampling to instances in which C-mode time-courses exceeded 1 standard deviation (**Figure S7**).

To further explore the temporal sequencing of C-mode occurrences and transitions (**Figure 5**), we defined for each dataset, a concatenated sequence of C-mode occurrences across subjects/animals, and calculated the transition probability matrix from a C-mode *i* at time *t* to another C-mode *j* at time *t*+ 1 as the proportion of transitions *i* → *j* and all other transitions from *i*^19,30^. Only transitions within the same subject were included, and we first considered sequences in which we counted the auto-transitions (*i* = *j*), namely persistence probabilities. Off-diagonal elements (*i* ≠ *j*), named transition probabilities were then computed after building sequences in which we removed the repeating elements in order to control for auto-correlations given the C-mode’s dwell time^19,79^. We quantified the directional prevalence of a transition (*P*_ij_ > *P*_ji_) by taking their difference. To measure the relative facilitation to reach a C-mode from another, we computed the Entropy of Markov Trajectories (HMT)^30,33^ from the off-diagonal matrix elements (i.e. transition probabilities). This method calculates the descriptive complexity of the paths between C-mode (in bits), where a lower complexity refers to less information required to access a destination C-mode *j* from a source C-mode *i*, hence being more accessible as the path travels through less path before reaching its destination. Specifically, for a Markov Chain (MC) defined by transition probability matrix P, we define the Entropy Rate per step *t* → *t* + 1 as:

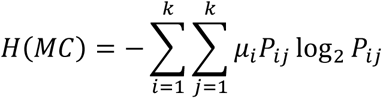

Where *μ* is the stationary distribution solving 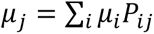. Also define the Entropy of a Markov Trajectory *T*_ij_ from C-mode *i* to C-mode *j* as:

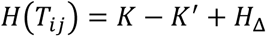

Where *K* = (*I* - *P* - *M*)^-1^(*H*^∗^ - *H*_Δ_), *M* is a matrix of stationary probabilities *M*_ij_ = *μ*_ij_; *I* the identity matrix; *H*^∗^ is the matrix of single-step entropies 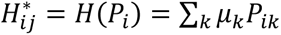 (from C-mode *i* to any C-mode *k*); and *H*_Δ_ is a diagonal entropy matrix with trajectories from a C-mode to itself (*H*_Δ_)*_ii_* = *H*(*MC*)/*μ*_i_, with zeros if *i* ≠ *j*^33^.

Statistical significance of persistence probabilities was tested from random sequences by generating 1000 permutations of C-mode occurrence sequences at the subject/animal level, concatenating the sequences and counting how many times a random iteration exceeded the true value to reach a p-value. Transition probabilities and directional transition prevalence were tested against matrices built after randomly permuting the non-repeating sequences 1000 times at the subject/animal level, concatenating them and removing samples if *Cmode*_i_(*t*+ 1) = *Cmode*_i_(*t*). These surrogates were also used to test the significance of Markov trajectory entropies. To further quantify and assess the relative complexity of transitions to a particular destination C-mode, we quantified, after repeating the analyses with subject level sequences, the sum of the HMT matrix columns, and compared their distributions across C-mode destinations (one-way ANOVA and Tukey test for multiple comparisons, **Fig 5C**). Finally, to assess the relationship of the mean GS-phase difference between occurrences of different C-modes and the HMT of their transitions, we computed the circular-linear correlations of their mean values (**Figure S7 and 5B** respectively) using the *circ_corrcl.m* Matlab function^78^.

### C-mode influence on static fMRI connectome and functional connectivity gradients

We first generated C-mode co-fluctuation matrices by cross-multiplying the mean C-mode map with itself^43^, which yielded a voxel-wise representation of the co-activating (or co-deactivating) peaks of BOLD activity. We then computed the group-level mean Functional Connectivity matrices by computing the Pearson’s Correlation between the concatenated time-series of voxels in all subjects/animals in each group^80^, and computed the correlation between the vectorized upper triangular part of the FC matrix to the weighted (by occurrence rate) average of the C-mode Co-fluctuation matrices^30^ (**Fig S9A-B**). Voxels were ordered according to the RSN they belong to (see **Fig 1C**).

We further explored if the hierarchies that dominate FC are in accordance with the dynamic structure of C-modes. We first computed the gradients of FC matrices from each species using the Diffusion Mapping method (BrainSpace^81^ - *diffusion_mapping.m* Matlab function: FC matrix sparse to top 10-percentile per node, 5 components, anisotropic diffusion parameter = 0.5)^76,82^. We mapped the gradients (**Fig S10**) and computed their spatial correlation to the obtained C-modes, matching each C-mode to the gradient with the highest absolute Pearson’s correlation (**Fig 6A**). Finally we plotted the variance explained by each gradient (lambda) to the group mean occurrence rate of their matched C-mode, fitting a linear model to this relationship (**Fig 6B**).

## ACKNOWLEDGMENTS

This work has received funding from the European Research Council (ERC) under the European Union’s Horizon 2020 research and innovation program (#DISCONN; no. 802371 to A.G.). The authors also acknowledge funding from the Simons Foundation (SFARI 982347 to A.G. and S.P).

## AUTHOR CONTRIBUTIONS

A.G. and D.G.B. conceived the study. D.G.B., A.G. and T.X. curated and preprocessed the data, and D.G.B. developed the computational analyses with input from A.G., T.X. and S.P. A.G. and D.G.B. wrote the manuscript with input from T.X., S.P., and B.S.R.

## DECLARATION OF INTERESTS

The authors declare no competing interests.

## DATA AVAILABILITY

All raw rsfMRI data can be found in the following repositories:

Human – HNU^57^: http://fcon_1000.projects.nitrc.org/indi/CoRR/

Human – MSC^58^: https://openneuro.org/datasets/ds000224/versions/1.0.3/download

Macaque – NC^61^: http://fcon_1000.projects.nitrc.org/indi/PRIME/newcastle.html

Mouse: Data will be made available upon publication of this work

## Notes

### Competing Interest Statement

The authors have declared no competing interest.

### Summary of Updates

In this version we have included some new analysis and fixed typos

