## Supplementary Figures for "Evolutionarily conserved fMRI network dynamics in the mouse, macaque, and human brain"

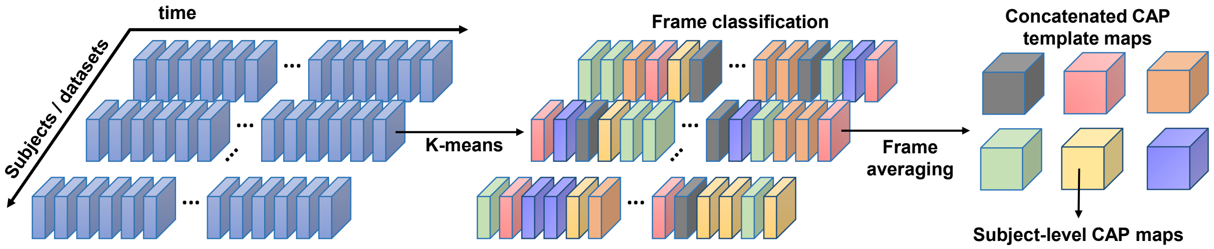


Figure S1. Data clustering and CAP detection (A) fMRI time-series were concatenated and fMRI frames were classified given their spatial similarity (Pearson's Correlation). Frames in a cluster were then averaged at the voxel level to create group-level CAP maps and T-score normalized. Subject-level CAP maps were then computed by averaging fMRI frames for each CAP.

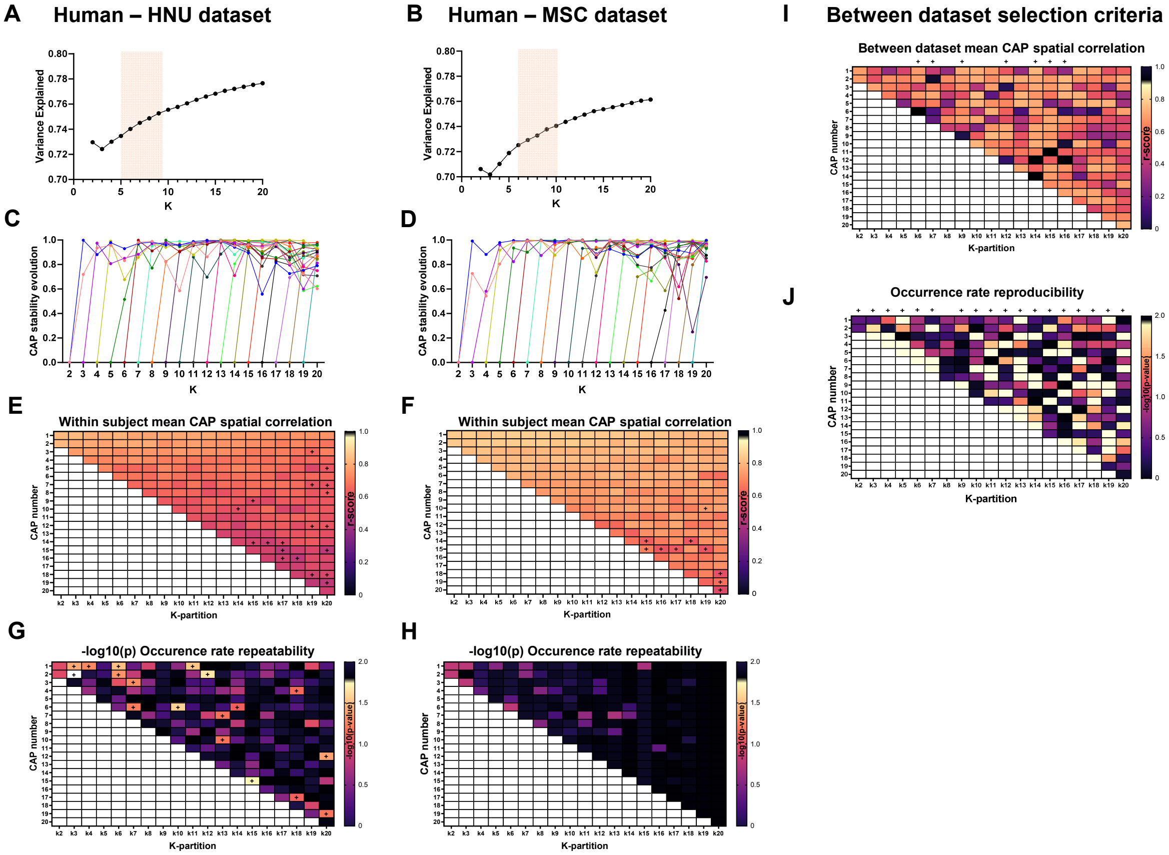


Figure S2. Clustering partition selection – human fMRI time-series. (A-B) Variance explained by clustering each independent dataset (A: HNU, and B: MSC) fMRI frames with k=2:20 (mean +/- SD for 5 runs). (C-D) Evolution of CAP stability, i.e, its ability to consistently be detected with the same anatomical configurations at higher partitions. For each partition (k) starting with k=3, the spatial correlation between the matched CAPs from k-1 (Hungarian Algorithm) is plotted, and a the new CAP at zero. (E-F) Within subject test-retest repeatability computed as the average spatial correlation between single-subject CAP-maps from all sessions. Asterisks denote CAPs within a partition (k) in which the within-subject correlation values was below a critical value of the highest correlation obtained from 1000 permutations with randomly shuffled fMRI frames, preserving occurrence rate. (G-H) Between session occurrence rate repeatability, computed as the –log(10) of the p-value of a Kruskal-Wallis test comparing the medians of all sessions. Asterisks denote CAPs within a partition (k) which showed a significant difference between sessions ($p > 0.05$). (I) Between dataset (HNU vs MSC) mean CAP spatial correlations. Asterisks denote partitions (k) in which at least one CAP was not significantly similar between datasets (1000 random permutations of frame reshuffling, preserving CAP occurrence rate). (J) Between dataset occurrence rate comparisons. Asterisks denote partitions (k) in which at least one matched CAP had significant differences in the distributions of its occurrence rate (Wilkoxon Signed Rank test, $p < 0.05$, FDR corrected for k comparisons).

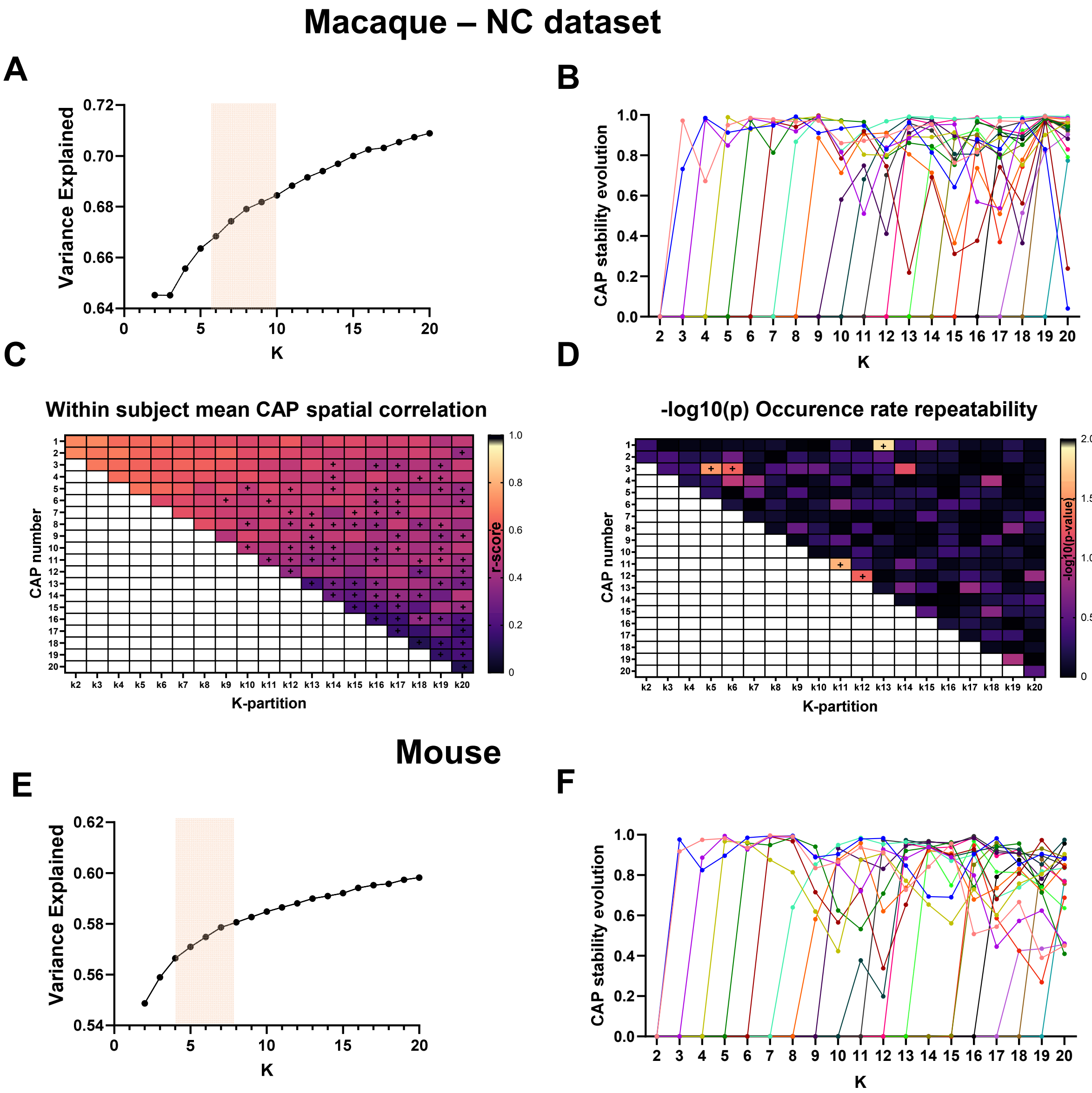


Figure S3. Clustering partition selection –Macaque and mouse fMRI time-series. (A) Variance explained by clustering the concatenated dataset with k=2:20 (mean +/- SD for 5 runs). (B) Evolution of CAP stability, i.e., its ability to consistently be detected with the same anatomical configurations at higher partitions. For each partition (k) starting with k=3, the spatial correlation between the matched CAPs from k-1 (Hungarian Algorithm) is plotted, and the new CAP at zero. (C) Within subject test-retest repeatability computed as the average spatial correlation between single-subject CAP-maps from all sessions. Asterisks denote CAPs within a partition (k) in which the within-subject correlation values was below a critical value of the highest correlation obtained from 1000 permutations with randomly shuffled fMRI frames, preserving CAP identity and occurrence rate. (D) Between session occurrence rate repeatability, computed as the –log(10) of the p-value of a Wilcoxon Signed Rank test comparing the medians of both sessions. Asterisks denote CAPs within a partition (k) which showed a significant difference between sessions (p > 0.05). (E) Variance explained by clustering the concatenated mouse dataset with k=2:20 (mean +/- SD for 5 runs). (F) Evolution of mice CAP stability, i.e., its ability to consistently be detected with the same anatomical configurations at higher partitions.

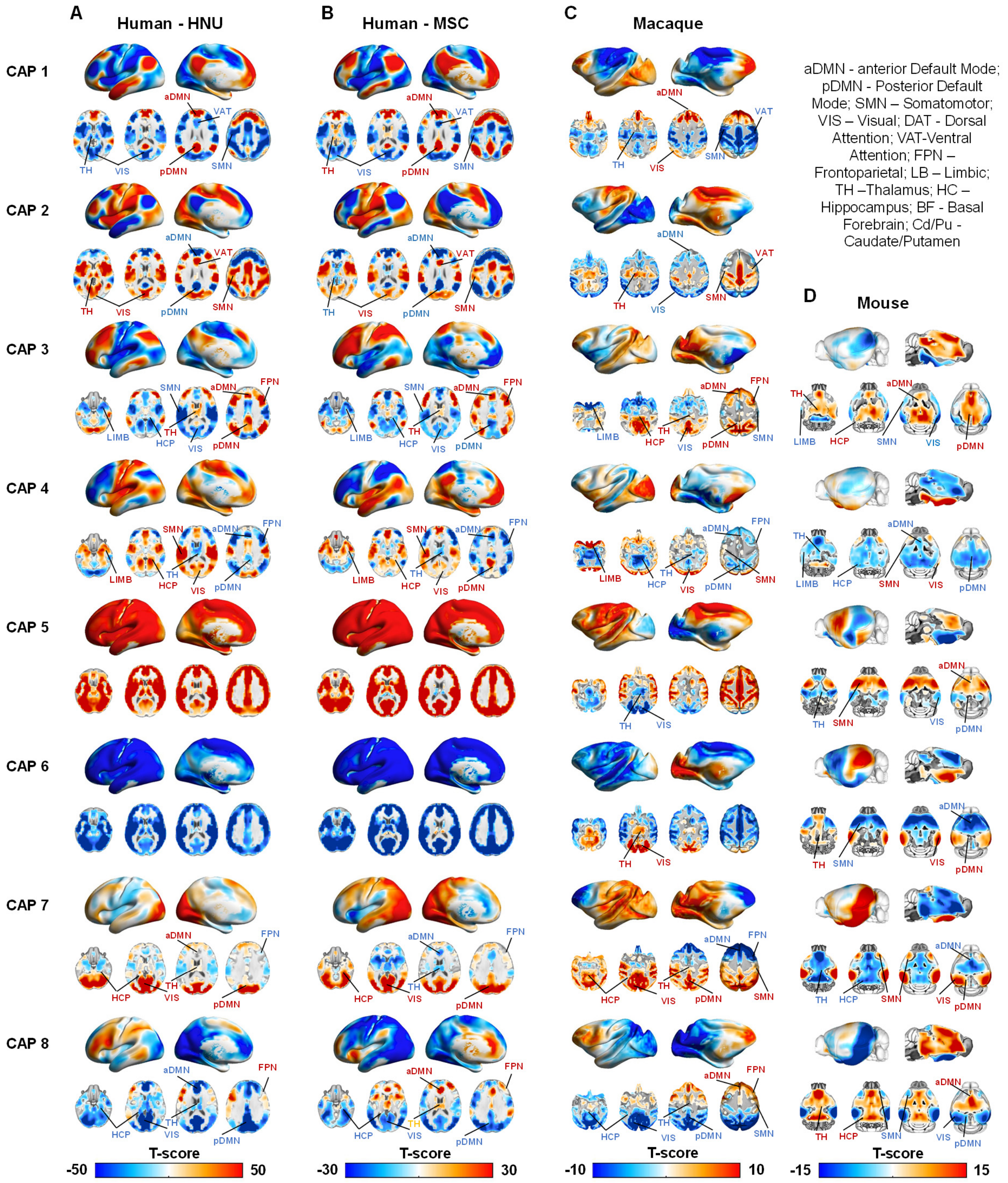


Figure S4. CAPs in awake humans, macaques and mice (A-B) CAPs (T-scores, $p < 0.05$ FDR corrected) of human datasets HNU (A) and MSC (B). (C) Macaque CAPs (T-scores, $p < 0.05$ FDR corrected). (D) Mouse CAPs (T-scores, $p < 0.05$ FDR corrected). Abbreviations: aDMN - anterior Default Mode; pDMN - Posterior Default Mode; SMN – Somatomotor; VIS – Visual; DAT - Dorsal Attention; VAT-Ventral Attention; FPN – Frontoparietal; LB – Limbic; TH –Thalamus; HC – Hippocampus; BF - Basal Forebrain; Cd/Pu - Caudate/Putamen.

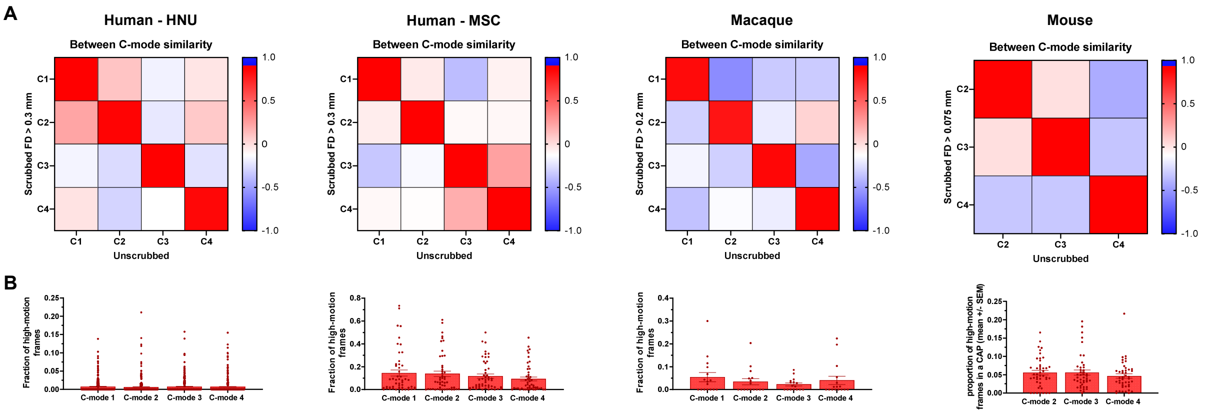


Figure S5. C-modes are not affected by high-motion frames (A) Spatial similarity matrices between C-modes obtained using scrubbed and non-scrubbed data of both human datasets (left), macaques (middle) and mice (right). (B) Distribution of the proportion of the fMRI frames within a C-mode that were flagged as high-head motion (FD > 0.3, 0.3, and 0.075 mm for humans, macaques, and mice respectively). No significant differences between C-modes were observed in all species/datasets (one-way ANOVA, $p > 0.1$ for all comparisons, uncorrected).

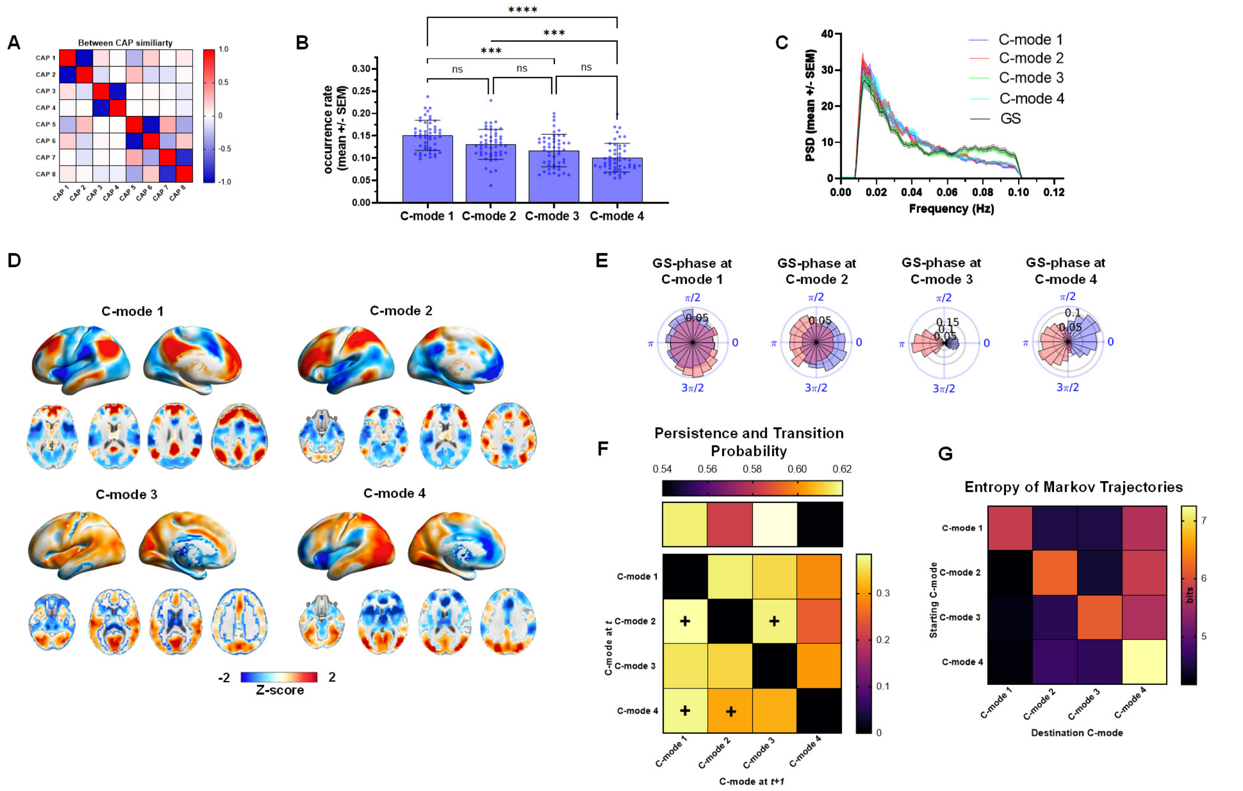


Figure S6. C-modes in MSC dataset reproduce findings from human HNU cohort. (A) Between CAP spatial similarity matrix confirming the existence of CAP-anti-CAP configurations. (B) Occurrence rates distributions of C-modes, ordered to their matched C-mode in the HNU dataset using the Hungarian Algorithm. (C) Group-level power spectral density (mean +/- SEM) of C-mode time-series, as well as for the Global fMRI Signal (GS). (C) C-mode maps from the subtraction of CAP-anti-CAP pairs divided by two. (E) GS-phase distributions at the occurrence of each C-mode, conditioned to the C-mode to frame correlation time-course to be > 1SD. Blue and red distributions correspond to GS phases sampled from the positive and negative C-mode time courses, respectively. All distributions significantly deviate from circular uniformity (Rayleigh test, $p < 0.05$, FDR corrected). (F) Persistence (top row) and transition (off-diagonal) probability. Red crosses denote the destination C-mode of preferred directional transitions ($P_{ij} > P_{ji}$). (G) Entropy of Markov trajectories (HMT) show that the C-modes with the highest accessibility are the ones from C-modes with higher occurrence rates. Higher entropy indicates lower accessibility of a destination C-mode (column) from a starting C-mode (row).

### GS-phase difference between C-mode ocurrences in a GS cycle

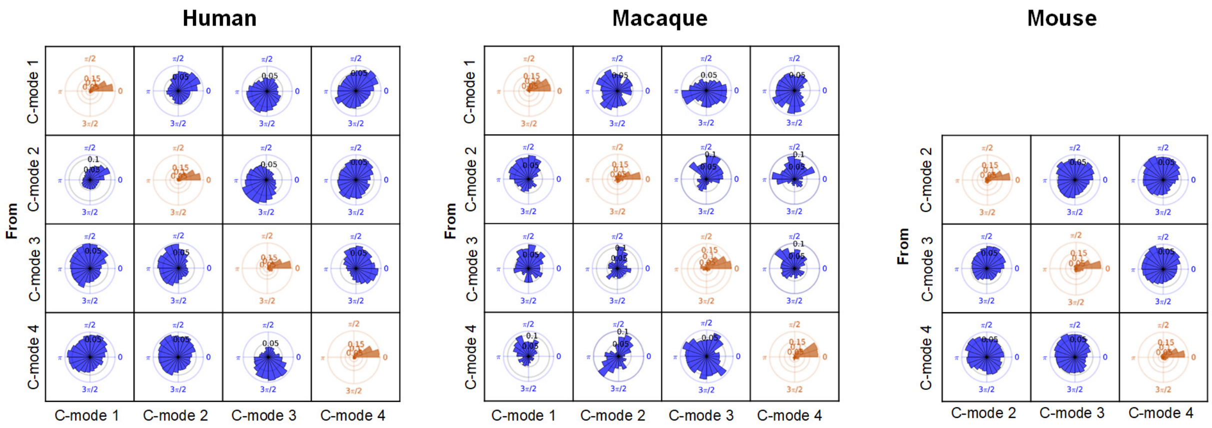


Figure S7. C-mode occurrences are phase-coupled within GS-cycles. Each panel corresponds to the circular distribution of GS phase differences between occurrences of a C-mode within a GS-cycle (rows), and the following occurrences of another C-mode (columns) within the same cycle or in an immediately subsequent one. Sampling was conditioned to occurrences in which the C-mode time-course is above 1SD. All distributions significantly deviate from circular uniformity ($p < 0.05$, Rayleigh test).

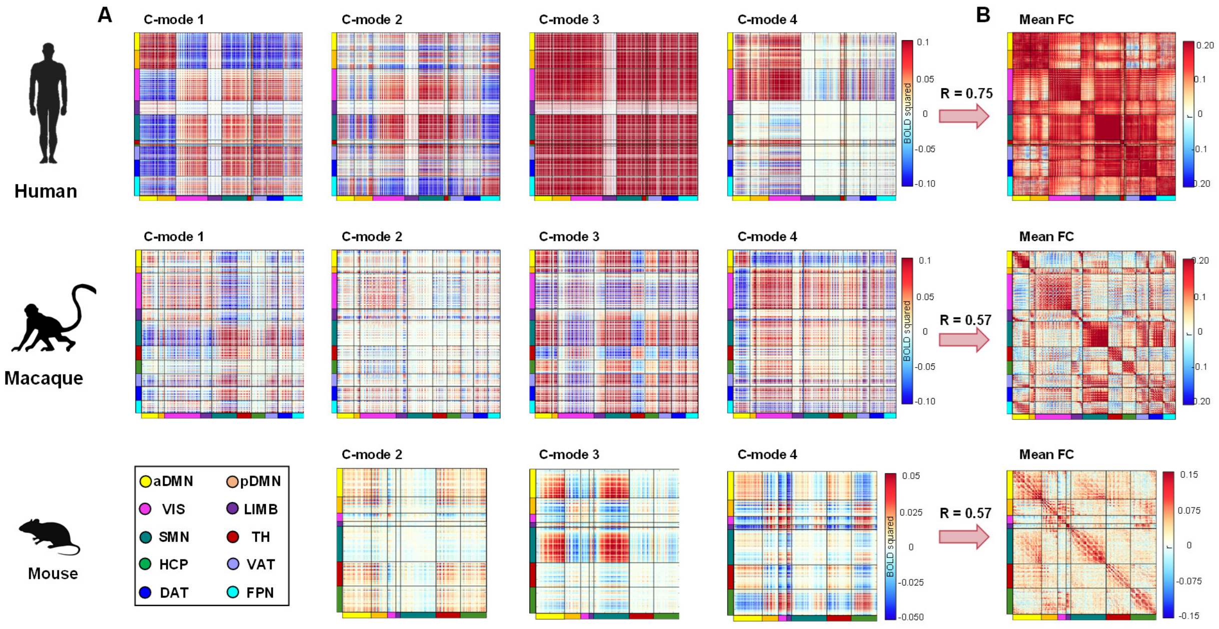


Figure S8. C-Modes topography and occurrence explain functional organization of static fMRI connectome. (A) C-mode Cofluctuation matrices obtain from the tensorial product the each C-mode mean map. (B) Group-level voxel-wise Functional Connectivity matrices (Pearson's correlation between concatenated time-series of voxels in all subjects/animals). The black arrow denotes the spatial correlation between vectorized form of the upper-triangular part of the mean FC matrix, and the weighted average of the triangular-parts of the C-modes' cofluctuation matrices (weighted by their average occurrence rates).

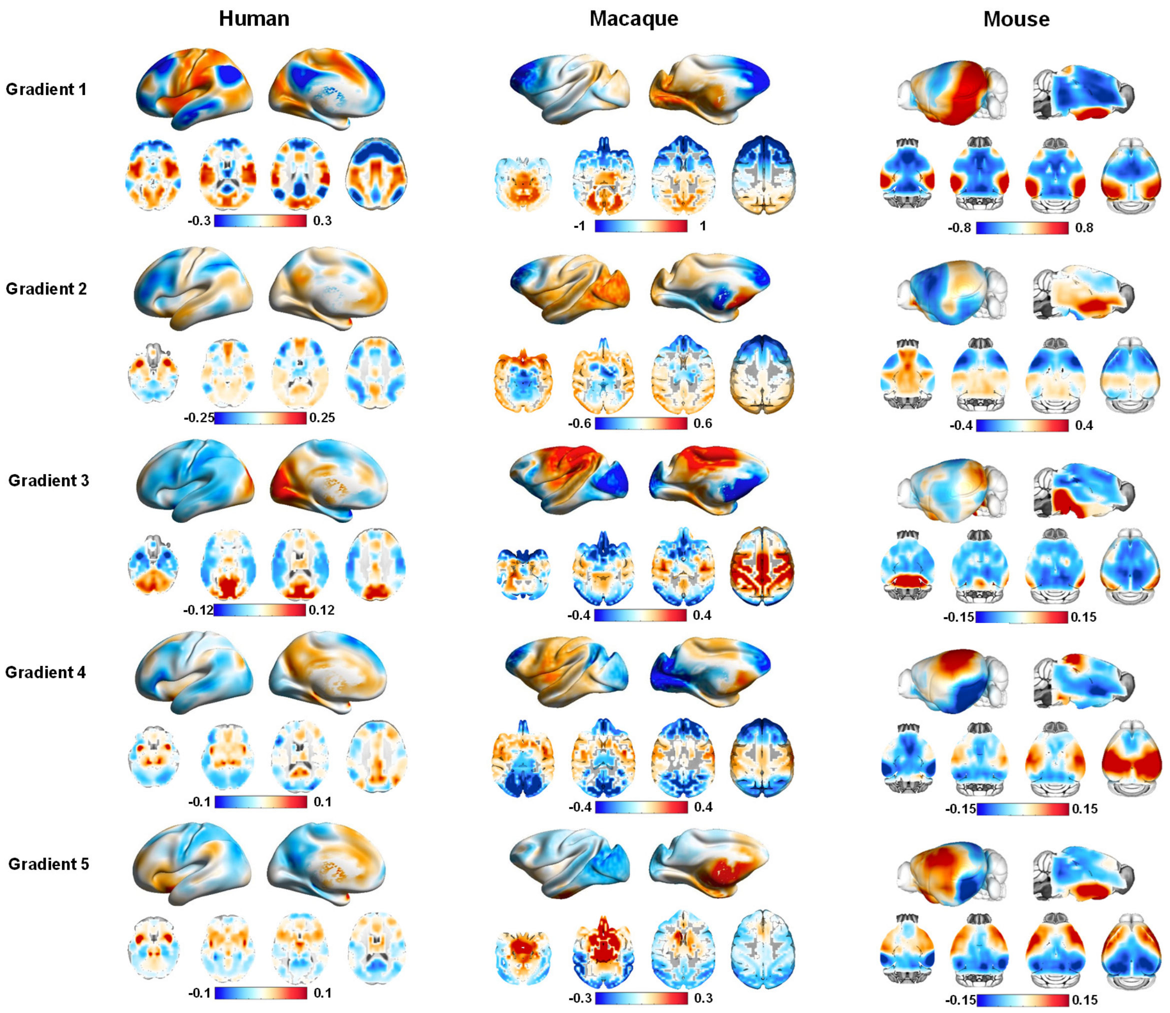


**Figure S9. Functional Connectivity gradients in awake humans, macaques, and mice.** Gradient maps are organized in order of explained connectome variance.
